# New cycle, same old mistakes? Overlapping vs. discrete generations in long-term recurrent selection

**DOI:** 10.1101/2021.10.12.464059

**Authors:** Marlee R. Labroo, Jessica E. Rutkoski

**Author notes:** E-mail addresses: ML JR.

## Abstract

**Background:** Recurrent selection is a foundational breeding method for quantitative trait improvement. It typically features rapid breeding cycles that can lead to high rates of genetic gain. In recurrent phenotypic selection, generations do not overlap, which means that breeding candidates are evaluated and considered for selection for only one cycle. With recurrent genomic selection, candidates can be evaluated based on genomic estimated breeding values indefinitely, therefore facilitating overlapping generations. Candidates with true high breeding values that were discarded in one cycle due to underestimation of breeding value could be identified and selected in subsequent cycles. The consequences of allowing generations to overlap in recurrent selection are unknown. We assessed whether maintaining overlapping and discrete generations led to differences in genetic gain for phenotypic, genomic truncation, and genomic optimum contribution recurrent selection by simulation of traits with various heritabilities and genetic architectures across fifty breeding cycles. We also assessed differences of overlapping and discrete generations in a conventional breeding scheme with multiple stages and cohorts.

**Results:** With phenotypic selection, overlapping generations led to decreased genetic gain compared to discrete generations due to increased selection error bias. Selected individuals, which were in the upper tail of the distribution of phenotypic values, tended to also have high absolute error relative to their true breeding value compared to the overall population. Without repeated phenotyping, these individuals erroneously believed to have high value were repeatedly selected across cycles, leading to decreased genetic gain. With genomic truncation selection, overlapping and discrete generations performed similarly as updating breeding values precluded repeatedly selecting individuals with inaccurately high estimates of breeding values in subsequent cycles. Overlapping generations did not outperform discrete generations in the presence of a positive genetic trend with genomic truncation selection, as past generations had lower mean genetic values than the current generation of selection candidates. With genomic optimum contribution selection, overlapping and discrete generations performed similarly, but overlapping generations slightly outperformed discrete generations in the long term if the targeted inbreeding rate was extremely low.

**Conclusions:** Maintaining discrete generations in recurrent phenotypic selection leads to increased genetic gain, especially at low heritabilities, by preventing selection error bias. With genomic truncation selection and genomic optimum contribution selection, genetic gain does not differ between discrete and overlapping generations assuming non-genetic effects are not present. Overlapping generations may increase genetic gain in the long term with very low targeted rates of inbreeding in genomic optimum contribution selection.

## Background

Quantitative trait improvement is achieved by cyclically increasing mean genetic value of breeding populations via recurrent selection. Recurrent phenotypic selection, reviewed by Hallauer & Darrah (1985), is a breeding strategy in which top-performing individuals are selected from a population and crossed to generate a new population for selection in the subsequent breeding cycle [1-3]. Recurrent phenotypic selection likely began with the invention of agriculture and is used to this day for quantitative trait improvement [3]. The advantage of this breeding strategy is that the breeding cycle length is short, as individuals can be selected as parents soon after they are born. Shorter cycle length leads to faster genetic gain, which is the rate of increase in mean genetic value due to selection in a population over time [4].

The main disadvantage of phenotypic selection is that selection accuracy tends to be low, because individuals are selected based on a single phenotypic observation, and selection accuracy directly impacts the rate of genetic gain [3]. This disadvantage is exacerbated at low trait heritabilities, as phenotypes are less indicative of true breeding values [5]. Different breeding schemes to improve the accuracy of phenotypic selection have been developed which involve testing families of progeny of selection candidates (e.g. half-sibs, full-sibs, or inbred lines) across multiple replicates or environments [3]. Most applied breeding programs of cereal crops are currently practicing some form of recurrent selection among families, especially inbred families. While selection by family improves accuracy, it also increases the breeding cycle length, which limits the rate of genetic gain that can be realized.

With the availability of genomic selection, recurrent selection schemes are being modified to use genomic estimated breeding values (GEBVs) rather than single phenotypic observations for parent selection [7-10]. This is often referred to as “rapid-cycle genomic selection” [11]. This approach can improve selection accuracy without increasing the breeding cycle length, thus increasing the rate of genetic gain. Recurrent phenotypic and genomic selection fundamentally differ in that estimates of breeding value based on phenotype are defined at the individual level, whereas GEBVs are defined at the marker or population level [7]. In recurrent phenotypic selection, individuals are phenotyped once prior to selection, and this comprises the only assessment of the individuals’ breeding values. In genomic selection, observations of marker effects or genetic relationships increase in number as new relatives are phenotyped. Thus, the accuracy of estimates of individual breeding values increases with genomic prediction even in absence of additional phenotypic data for evaluated individuals [7]. For example, an individual with a high true breeding value may have a low estimated breeding value in a given genomic selection cycle due to error, but in a subsequent cycle its breeding value estimate may be higher—in better agreement with its true breeding value—as the prediction model is updated with information from relatives.

This raises the question: if possible, should individuals from previous selection cycles be considered again as selection candidates in subsequent cycles? Or, in other words, should generations be allowed to overlap in phenotypic and genomic recurrent selection programs? Conventionally, individuals are only considered as candidates for selection during the cycle when they are evaluated. However, in clonally propagated or perennial species, individuals could be selected directly as parents for multiple seasons. In self-compatible species with multiple inflorescences, selected individuals could be self-pollinated and the resultant seed could be used for crossing in multiple selection cycles, even though the selfed progeny would not be identical to the parent genotype. In practice, it is common for plant breeders to recycle favored parents across cycles of selection, leading to overlap, even if the parent has not been phenotyped and statistically evaluated alongside the current selection candidates. The effect on genetic gain of maintaining discrete or overlapping selection generations has not been formally evaluated or reported. Given that selection accuracy may vary with cycle in breeding individuals from previous generations in genomic but not phenotypic selection, we hypothesized that allowing overlapping generations may be more favorable for rapid recurrent genomic selection compared to rapid recurrent phenotypic selection. Unexpectedly, we found that overlapping generations decreased the rate of genetic gain under phenotypic selection compared to discrete generations.

This study had two primary objectives: 1) to determine if generations should be overlapping or discrete in phenotypic and genomic recurrent selection programs, and 2) to determine in what selection scenarios overlapping and discrete generations can be recommended for recurrent selection. The effects of overlapping and discrete generations on the inbreeding rate, average parental age, and the selection accuracy were also examined.

## Methods

Stochastic simulations in the R package AlphaSimR were conducted to examine various recurrent selection scenarios [12]. All simulations were run on the Biocluster High Performance Computing system housed in the Carl R. Woese Institute for Genomic Biology at the University of Illinois at Urbana-Champaign and maintained by the Computer Network Resource Group. Two main trait and pipeline architectures were considered: 1) recurrent selection on a purely additive trait in a single cohort per breeding cycle (RS-A), and 2) recurrent selection on a trait with additive, year, and additive x year effects with multiple cohorts per breeding cycle (RS-AY). For both architectures, an outbred, diploid, hermaphroditic founder population was generated with the *runMacs* function. Individuals had ten chromosomes with 1,000 segregating sites per chromosome.

For the RS-A scenarios, with the purely additive trait, 100 sites per chromosome were assigned additive effects and 50 sites per chromosome were genotyped by a simulated SNP-chip. Additional File 1 contains the script used to generate the base founder population. To start each simulation replicate, 100 individuals were drawn from the founder population. Starting mean genetic value was 0, genetic variance was 1, error variance was 4, and narrow-sense heritability was either 0.1, 0.5, or 0.9. In the first year, 20 parents were selected phenotypically. See Additional File 2 for the script used to start each simulation. After the first year, a breeding cycle consisted of crossing the selected parents, phenotypic evaluation and parent selection before flowering, then restarting the cycle by making 100 random crosses of the selected parents which produced 1 progeny per cross (Fig. 1).

**Figure 1.**
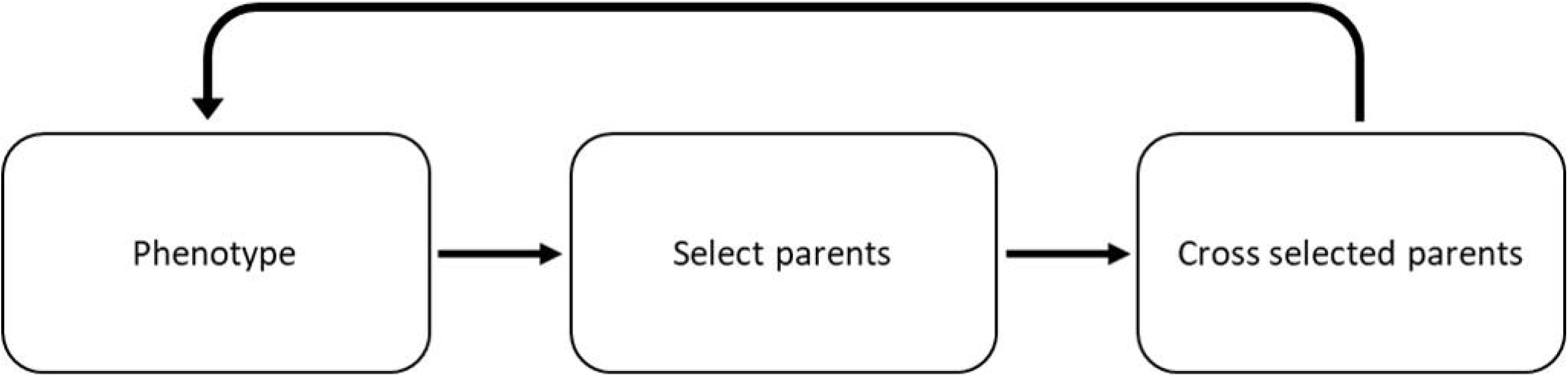
Overview of recurrent mass selection scheme for RS-A scenarios. For the RS-A scenarios, only the parental selection units varied in this study. For an overview of the RS-AY scenarios, see the Conventional scenario in Gaynor et al., 2017 [14].

Several factors were considered in the RS-A scenario (Fig. 2). Parents were selected from either discrete or overlapping generations. For discrete generations, parents were only selected from the current breeding cycle. For overlapping generations, parents were selected from any breeding cycle. Then, the selection on either phenotypic value, true genetic value, or GEBV as estimated by ridge regressed best linear unbiased prediction (RR-BLUP) was used. In phenotypic selection only, selection on either unreplicated phenotypes or thrice-replicated phenotypes was considered; in all other cases, phenotypes were unreplicated. In the case of genomic selection only, truncation vs. optimum contribution selection (OCS), as well as training the model on all generations (allGen) vs. training on the most recent previous five generations (fiveGen) to mimic what may occur in practical situations were also considered (Fig. 2). If selection occurred on phenotype or true genetic value, truncation selection of the top 20 individuals was always used. In the genomic selection scenarios, either truncation selection of the top 20 individuals was used or OCS was used with minimum effective population sizes (*N*_*e*_) of 10, 45, and 90. Higher minimum effective population size implied stricter control of inbreeding. OCS was implemented with the R package optiSel [13]. All RS-A scenarios were run for 50 breeding cycles and replicated across 10 simulations. See Additional File 1 for custom optiSel functions used in the study, and see Additional File 3 for the core script used to run the RS-A simulations.

**Figure 2.**
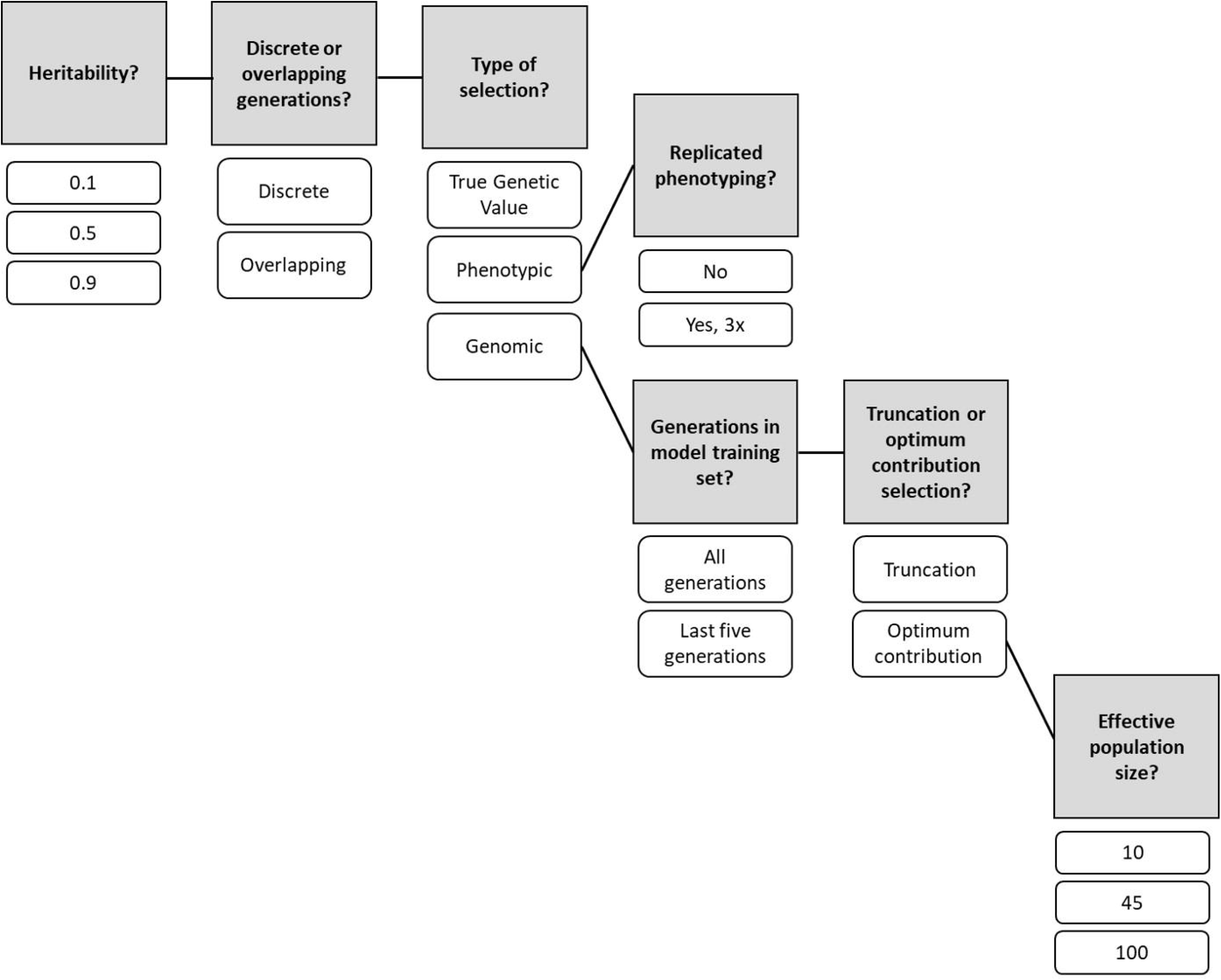
Overview of the RS-A scenario factors. Shaded boxes indicate factors and unshaded boxes indicate levels of factors. Solid lines connecting shaded boxes indicate that all combinations of factor levels were tested, while solid lines connecting unshaded factor levels to shaded factors indicate the subsequent shaded factors only apply to the connected factor level.

For the RS-AY scenario, with selection on an additive, year, and additive x year trait and multiple cohorts per cycle, a modification of the general breeding scheme of the Conventional Program described in Gaynor et al., 2017, was used [14]. As in the RS-A scenarios, 100 segregating sites per chromosome were assigned additive effects, and 50 sites per chromosome were genotyped by a simulated SNP-chip. To start each simulation replicate, 100 individuals were drawn from the founder population. Starting mean genetic value was 0, and genetic variance was 1. Additional File 4 contains the script used to start the RS-AY scenarios, and Additional File 5 contains a script to store the year effects. Phenotypes in subsequent stages were simulated using a custom R script according to the assumptions of a compound symmetry model. Phenotypes were not simulated with the Finlay-Wilkinson model, which is the default in AlphaSimR for traits with genotype x environment interactions. Year effects were drawn from a normal distribution with mean 0 and variance 0.2. Additive x year effects for each site were drawn from a normal distribution with mean 0 and variance scaled to achieve the targeted total additive x year variance of 0.2. As such, the variance of the distribution from which the additive x year effects were drawn was the variance of the additive marker effects times the targeted additive x year variance of 0.2 divided by the genetic variance of 1 in the base population. Plot error effects were drawn from a normal distribution with mean 0 and variance scaled to achieve variable broad-sense heritabilities at each stage in the breeding cycle, with heritability increasing at later stages compared to earlier stages. The heritabilities used differed from those in Gaynor et al., 2017 [14]. Phenotypes were the sum of the additive, year, additive x year, and plot error effects.

For the RS-AY scenario, 30 selected parents entered the breeding pipeline at stage 1 and were crossed randomly into 100 biparental crosses with 97 progeny each. In stage 2, doubled haploid lines were produced from each of the year 1 progeny. In stage 3, the doubled haploid lines were phenotyped in headrows from which 500 individuals are advanced. In stage 4, the 500 individuals advanced from stage 3 were then phenotyped in a preliminary yield trial, and 50 individuals were advanced. In stage 5, the 50 individuals advanced from stage 4 entered an advanced yield trial, from which 10 individuals were advanced. In stage 6, the 10 individuals advanced from stage 5 were phenotyped in an elite yield trial, and all individuals were advanced. In stage 7, all individuals from stage 6 were reevaluated in the second year of the elite yield trial. In stage 8, a single variety was chosen from the elite yield trials. In RS-AY scenarios with discrete generations, the 20 top-ranked individuals from stage 4 and all individuals from stage 5 of the most recent cycle were selected as parents (modified from Gaynor et al., 2017, in which the crossing block was composed of the 20 top-ranked individuals from stage 4, the 10 top-ranked individuals from stage 5, and also the 20 best individuals from the crossing block of the previous cycle, which implicitly allowed overlapping generations) [14]. In scenarios with overlapping generations, the 20 top-ranked individuals from stage 4 and the 10 top-ranked individuals from stage 5 were selected as parents from all cycles conducted in the breeding program. In the genomic selection scenarios, all records from stages 4-7 from all cycles conducted in the breeding program comprised the training set, regardless of whether generations were overlapping or discrete. Each stage was assumed to take one year. The breeding program was run for 40 years. The scripts to run each RS-AY scenario are located in Additional Files 6—9.

For each parent selection scenario in RS-A, mean genetic value was always recorded in the current generation of individuals in a given cycle to examine the genetic trend due to selection. For RS-AY, mean genetic value was recorded in the current generation of parents in a given year. For both situations, selection error bias, mean genomic inbreeding, selection accuracy, and average parental age were also recorded in the selected parents of the current generation only. Selection error bias per cycle was the ratio of absolute error in the selected parents to absolute error in all selection candidates, where error was the deviation of the phenotype or GEBV from the true genetic value. For RS-AY, selection error bias was decomposed into component error due to year, additive x year, and plot error. The ratio of each absolute component error in the selected parents to absolute component error in all selection candidates was the selection error bias for the component. Mean genomic inbreeding per cycle was the average probability of allelic identity-by-descent between pairs of individuals, where identity-by-descent was tracked directly via the *setTrackRec()* option rather than estimated. Selection accuracy was Pearson’s correlation of GEBV or phenotype and the simulated true breeding value (TBV). By definition, selection accuracy was one for scenarios with selection on TBV. See Additional File 10 for the raw response variables from each simulation replicate and cycle (for RS-A) or year (for RS-AY).

To test for differences in responses by scenario for RS-A, time points representing the short-term, medium-term, and long-term were chosen as cycle 5, cycle 25, and cycle 45 respectively. For RS-AY, differences in responses were only interrogated at the terminal year 40. The RS-A and RS-AY scenarios were considered separate experiments. The RS-AY experiment was conceived subsequently to RS-A in order to explore additional sources of selection error bias (i.e. year and genotype x year effects).

For each time point, and for all responses studied except mean parental age and year error bias, the following linear model was constructed with the R package nlme:

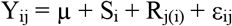

Y_ij_ was the response of interest for the i^th^ scenario and the j^th^ simulation replicate, µ was the grand mean, S_i_ was the fixed effect of the i^th^ scenario, R_j(i)_ was the random effect of the j^th^ simulation nested in the i^th^ scenario with N(0, σ_j(i)_ ^2^), and ε_ij_ was the random residual error with N(0, Rσ_ε_ ^2^) where σ_ε_ ^2^ was the error variance, and R was a matrix whose diagonal was a weighting factor used to model unique error variances for each scenario [15]. Differences in means by scenario were tested by the *anova*.*lme* function in nlme [15]. Pre-planned contrasts of differences in responses by scenario were made at α = 0.05 with the *pairs* function in the R packages emmeans for the discrete vs. overlapping variations of otherwise identical scenarios [16, 17]. Contrasts for OCS at *N*_*e*_ = 10 were not possible in the long term because the optimization of GEBV and mean genomic inbreeding ceased to solve around cycle 35.

Because mean parental age in the selected individuals was uniformly one with no variance in the RS-A discrete scenarios, Student’s *t* test was conducted with the *t*.*test* function in R to test whether mean parental age at each timepoint significantly differed from μ = 1 for each overlapping scenario at α = 0.05 subject to Bonferroni correction given the number of tests in the family. Because mean parental age for the RS-AY discrete scenarios was uniformly 3.67, Student’s *t* test was conducted as above to test whether mean parental age significantly differed from μ = 3.67. Similarly, because year error bias was uniformly one with no variance in the RS-AY discrete scenarios, Student’s *t* test was used to examine whether mean year error bias significantly differed from μ = 1 for the RS-AY overlapping scenarios at α = 0.05 subject to Bonferroni correction given the number of tests in the family. (Year error bias was 1 in the discrete scenarios because all candidates were evaluated in the same year and therefore had the same year value.)

## Results

### Genetic trends

In the RS-A case, significant differences in mean genetic value by scenario were observed (Additional File 11). In terms of mean genetic value, unreplicated discrete phenotypic selection outperformed unreplicated overlapping phenotypic selection in the long term for all heritabilities, and in the medium term if *h*^*2*^ = 0.1 or 0.5 (Fig. 3; Additional File 12). Performance of unreplicated discrete and overlapping phenotypic selection did not significantly differ in the short term (Fig. 3; Additional File 12). If phenotyping was replicated three times, then discrete phenotypic selection outperformed overlapping in the long and medium term if *h*^*2*^ = 0.1 or 0.5, and in the short term if *h*^*2*^ = 0.1 only (Fig. 3; Additional File 12). In contrast, if true genetic value was used for selection, then mean genetic value of discrete vs. overlapped selection did not differ significantly at any timepoint.

**Figure 3.**
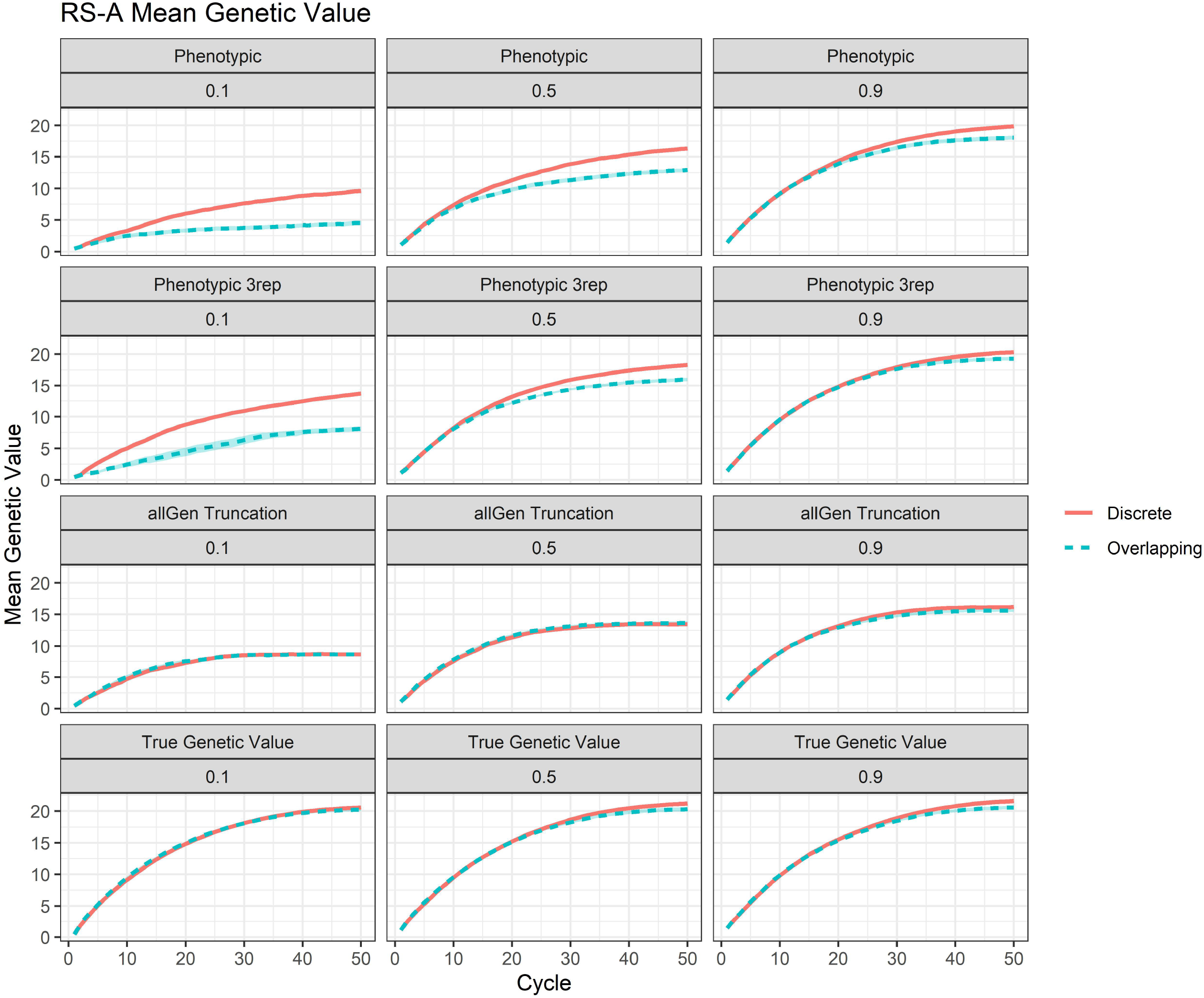
Mean genetic value for selected RS-A scenarios. Mean genetic value per cycle for the RS-A scenarios of phenotypic selection, thrice-replicated phenotypic selection, genomic truncation selection with all generations used in the training set (allGen truncation), and selection on true genetic value. Values are surrounded by the 95% confidence interval of the cycle mean.

Discrete and overlapping generations appeared to perform similarly with genomic selection in the RS-A scenarios (Fig. 3; Additional File 12—13). The exceptions were that overlapping generations always outperformed discrete generations with OCS at *N*_*e*_ = 100 and *h*^*2*^ = 0.5 or 0.9 regardless of training set used, and in the long term discrete generations outperformed overlapping with OCS at *N*_*e*_ = 45 and *h*^*2*^ = 0.9 with training on the previous five generations (Additional File 12—13). Also, in the short term, overlapping generations outperformed discrete with OCS at *N*_*e*_ = 100 at *h*^*2*^ = 0.5 or 0.9 with training on the previous five generations as well as training on all generations (Additional File 12—13).

In the RS-AY case, significant differences in mean genetic value by scenario were observed at year 40 (Additional File 11). Discrete genomic selection outperformed overlapping genomic selection, and discrete phenotypic selection outperformed overlapping phenotypic selection (Fig. 4; Additional File 12).

**Figure 4.**
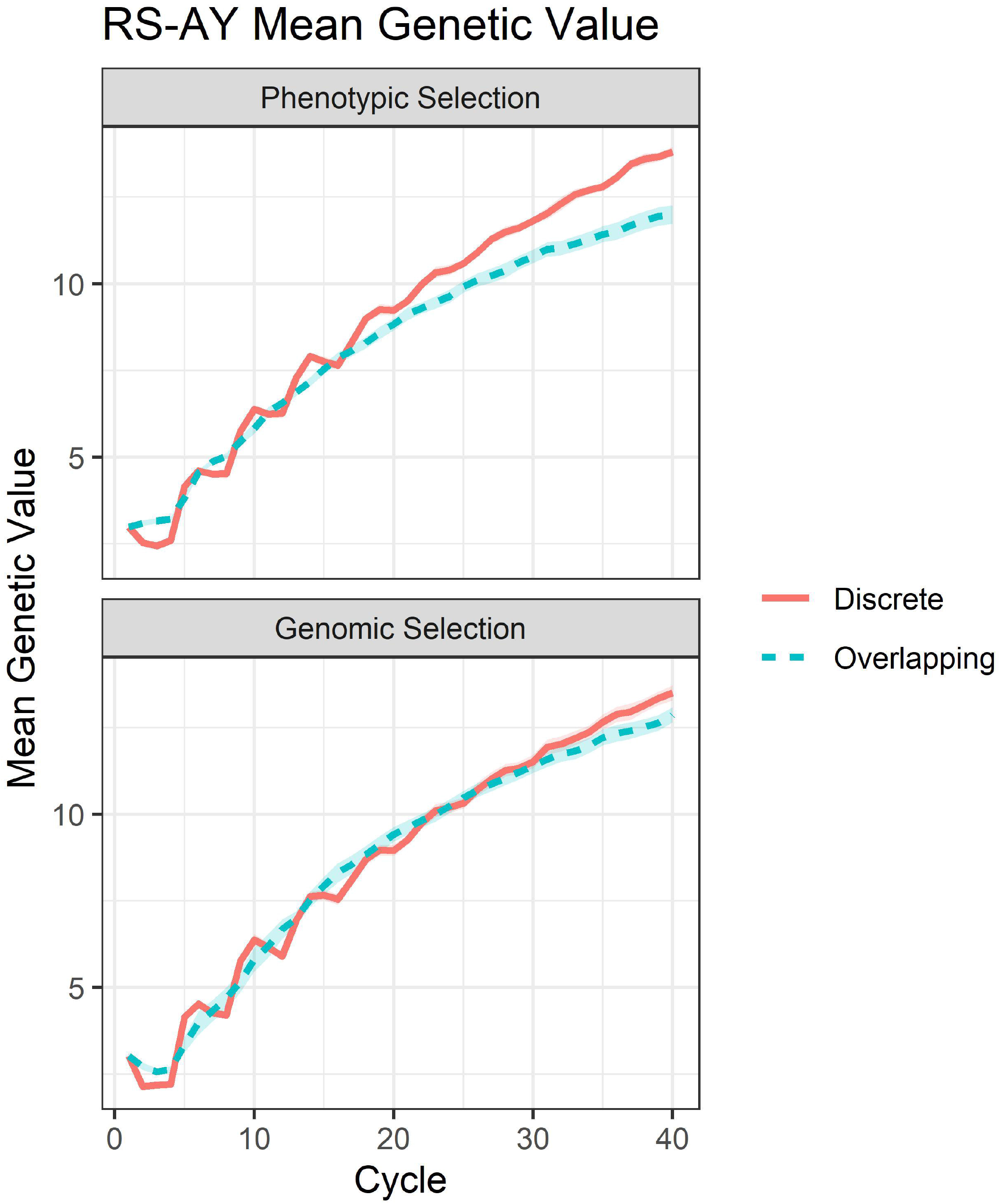
Mean genetic value for RS-AY scenarios. Mean genetic value per cycle for the RS-AY scenarios of phenotypic selection and genomic selection surrounded by the 95% confidence interval of the cycle mean.

### Selection error bias

For the RS-A cases, significant differences in mean selection error bias by scenario were observed (Additional File 11). For unreplicated phenotypic selection, selection error bias was always higher in overlapping selection scenarios, except in the short- and medium-term for *h*^*2*^ = 0.9 (Fig. 5; Additional File 12). Notably, this pattern mirrors the observed trend in mean genetic value. If phenotyping was replicated three times, selection error bias remained higher in overlapping generations in the same scenarios as unreplicated phenotypic selection (Fig. 5; Additional File 12). With selection on true genetic value, by definition selection error bias did not differ between overlapping and discrete generations, as error for all candidates was zero (Fig. 5). For genomic truncation selection, selection error bias also did not differ between overlapping and discrete scenarios at any point if the training set was composed of all generations (Fig. 5; Additional File 12). However, if the training set was composed of the previous five generations, then selection error bias in overlapping scenarios was significantly higher than discrete in the long-term with genomic truncation selection (Additional File 12, 14).

**Figure 5.**
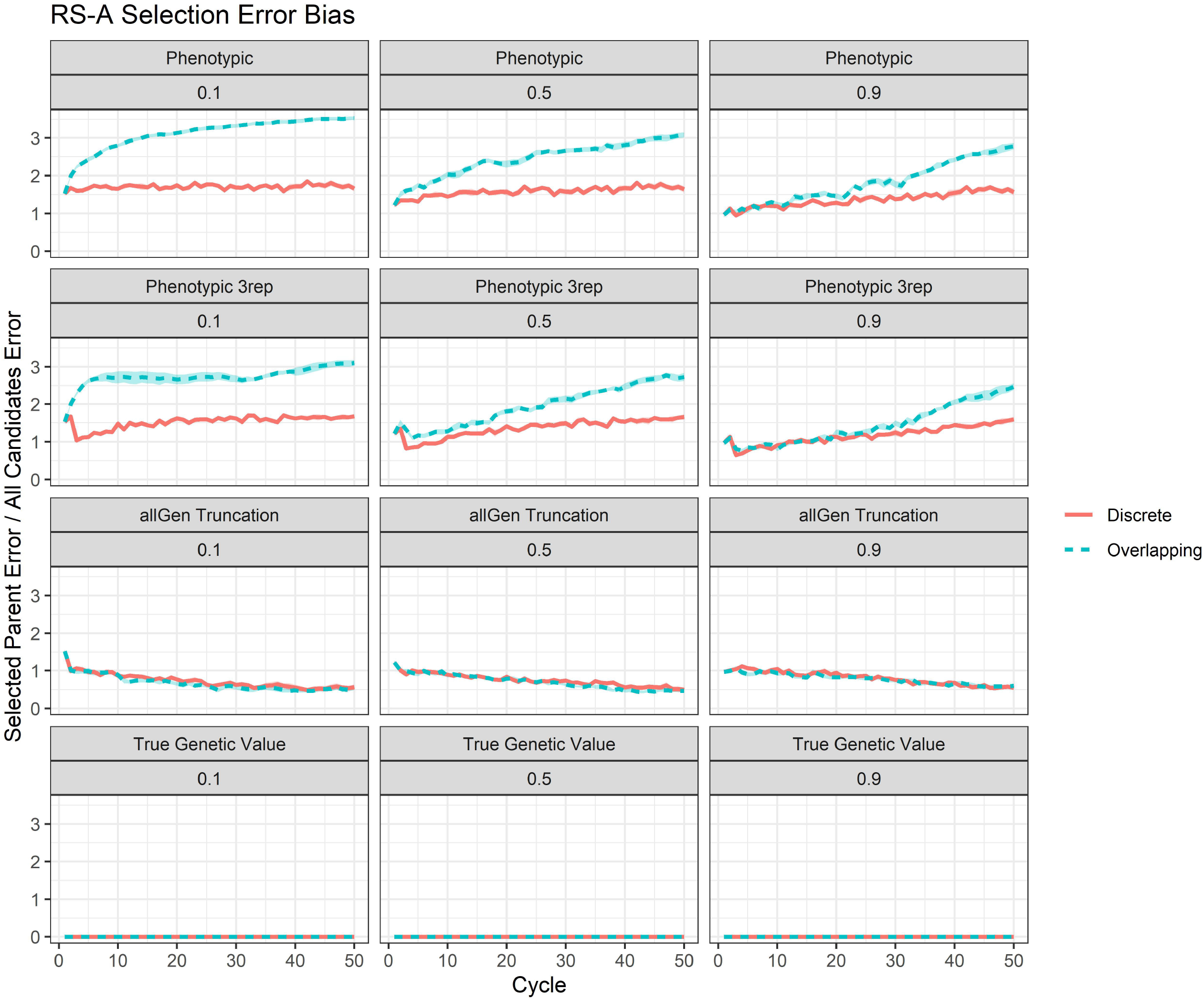
Selection error bias for selected RS-A scenarios. Selection error bias per cycle for the RS-A scenarios of phenotypic selection, thrice-replicated phenotypic selection, genomic truncation selection with all generations used in the training set (allGen truncation), and selection on true genetic value. Values are surrounded by the 95% confidence interval of the cycle mean.

For genomic OCS with the training set composed of all generations, discrete and overlapping selection error bias did not significantly differ except in the short and medium term if *N*_*e*_ = 100 and *h*^*2*^ = 0.5 or 0.9, in which case overlapping selection error bias was significantly higher (Additional File 12, 14). If the training set was composed of the previous five generations, then in the short term selection error bias did not significantly differ except at *N*_*e*_ = 100 for *h*^*2*^ = 0.5 or 0.9, in which case overlapping selection had a higher selection error bias (Additional File 12, 14). In the medium and long term with training on the previous five generations, discrete always had higher selection error bias than overlapping (Additional File 12, 14).

For the RS-AY cases, significant differences in mean selection error bias by scenario were observed (Additional File 11). Discrete phenotypic selection had significantly lower selection error bias than overlapping phenotypic selection, but no significant difference was observed for discrete vs. overlapping genomic selection (Fig. 6; Additional File 12). Significant differences in additive x year error bias and plot error bias were also observed (Fig. 6; Additional File 11).

**Figure 6.**
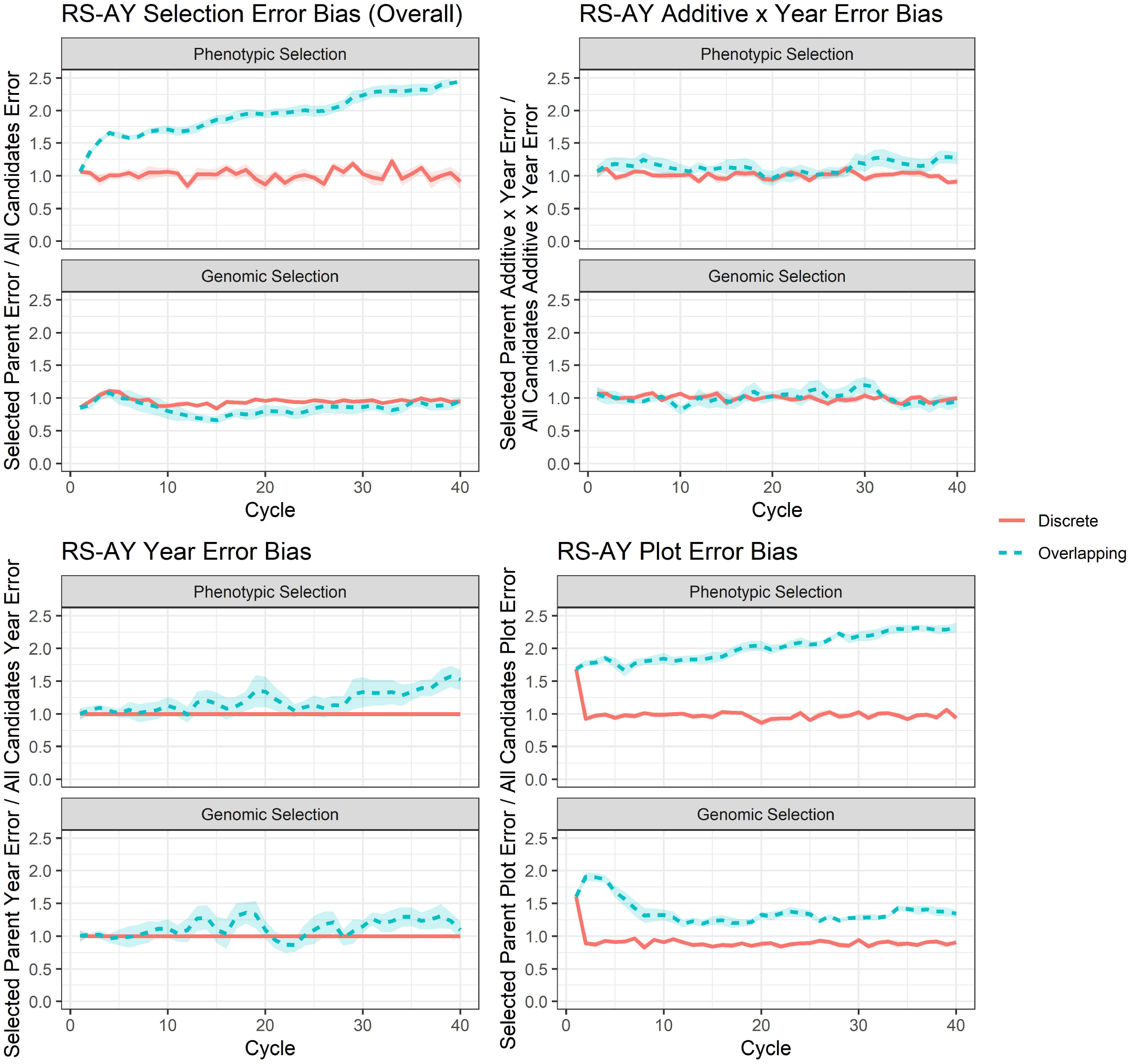
Selection error bias for RS-AY scenarios. Selection error bias per cycle for the RS-AY scenarios of phenotypic selection and genomic selection surrounded by the 95% confidence interval of the cycle mean. Overall selection error bias is show as well as error bias due to year, additive x year, and plot error.

Discrete phenotypic selection had significantly lower additive x year error bias than overlapping phenotypic selection, but no significant difference was observed for discrete vs. overlapping genomic selection (Fig. 6; Additional File 12). On the other hand, plot error bias was significantly lower for discrete vs. overlapping phenotypic selection and discrete vs. overlapping genomic selection (Fig. 6; Additional File 12). Year error bias significantly differed from 1 with overlapping phenotypic selection, but did not significantly differ from 1 with overlapping genomic selection (Fig. 6; Additional File 13).

### Mean genomic inbreeding

Significant differences in mean genomic inbreeding by scenario were observed in the RS-A cases (Additional File 11). For unreplicated and thrice-replicated phenotypic selection, mean genomic inbreeding was significantly higher with discrete selection at *h*^*2*^ = 0.1 at all time points but did not significantly differ for other heritabilites (Additional File 12, 15, 16). Mean genomic inbreeding did not significantly differ with selection on true genetic value (Additional File 12, 17). For genomic truncation selection with training on all generations, no significant differences in mean inbreeding were observed between discrete and overlapping scenarios except in the long term at *h*^*2*^ = 0.9, for which overlapping generations led to higher inbreeding than discrete (Additional File 12, 18). With training on the previous five generations, overlapping truncation genomic selection led to higher inbreeding in the short term at *h*^*2*^ = 0.1 and in the medium term at *h*^*2*^ = 0.5 and 0.9, but with no significant diffferences in the long term (Additional File 12, 19).

With genomic OCS, discrete selection sometimes led to higher inbreeding than overlapping selection despite optimization of the inbreeding rate. With training on all generations, this occurred for *h*^*2*^ = 0.1 in the medium term for *N*_*e*_ = 10 and the short and medium terms for *N*_*e*_ = 45, but did not occur for *N*_*e*_ = 100 (Additional File 12, 20—22). For *h*^*2*^ = 0.5, this occurred in the medium term for *N*_*e*_ = 10, and the medium and long term for *N*_*e*_ = 45 (Additional File 12, 20—21). However, in the short and medium term at *h*^*2*^ = 0.5 with training on all generations, overlapping led to higher inbreeding than discrete at *N*_*e*_ = 100 (Additional File 12, 22). For *h*^*2*^ = 0.9, discrete selection led to higher inbreeding in the short and medium term at *N*_*e*_ = 10, the medium term at *N*_*e*_ = 45, and the short term only at *N*_*e*_ = 100 (Additional File 12, 20—22).

With genomic OCS and training on the previous five generations, discrete selection led to higher rates of inbreeding in the medium and long term at *h*^*2*^ = 0.1 for all levels of *N*_*e*_, and additionally in the short term for *N*_*e*_ = 45 (Additional File 12, 23—25). At *h*^*2*^ = 0.5, discrete selection again led to higher inbreeding in the short term if *N*_*e*_ = 45 and the medium term for *N*_*e*_ = 45 and 100 only (Additional File 12, 24—25). At *h*^*2*^ = 0.9, discrete selection led to higher inbreeding rates in the short term for *N*_*e*_ = 10, lower inbreeding rates in the short term if *N*_*e*_ = 100, higher inbreeding rates in the medium and long term for *N*_*e*_ = 45, and higher inbreeding rates in the short and long term for *N*_*e*_ = 100 (Additional File 12, 23—25).

With RS-AY, significant differences in mean genomic inbreeding by scenario were also present at year 40 (Additional File 11). Discrete phenotypic selection led to significantly higher inbreeding than overlapping phenotypic selection, and discrete genomic selection also led to significantly higher inbreeding than overlapping genomic selection (Additional File 12, 26).

### Genetic variance

Significant differences in mean genetic variance by scenario were observed in RS-A (Additional File 11). For unreplicated phenotypic selection, significant differences in genetic variance in the current generation were only observed at *h*^*2*^ = 0.1 in the medium and long term, with overlapping selection maintaining higher genetic variance (Additional File 12, 15). For replicated phenotypic selection, genetic variance was significantly lower with overlapped selection only in the long term at *h*^*2*^ = 0.1 (Additional File 12, 16). No significant differences in genetic variance were observed for selection on true genetic value (Additional File 12, 17). For genomic truncation selection, no significant differences in genetic variance were observed regardless of training set or heritability (Additional File 12, 18—19).

For genomic OCS, no significant differences in genetic variance were observed if all generations were used in the training set (Additional File 12, 20—22). If the previous five generations were used in the training set, then at all heritabilities overlapping selection maintained greater genetic variance than discrete in the medium term if *N*_*e*_ = 100 only, while if *N*_*e*_ = 45 overlapping had higher genetic variance only if *h*^*2*^ = 0.5 or 0.9 (Additional File 12, 23—25). In the long term, overlapping selection maintained greater genetic variance if *N*_*e*_ = 45 at *h*^*2*^ = 0.1 or 0.9, and if *N*_*e*_ = 100 at all heritabilities (Additional File 12, 23—25).

For the RS-AY scenarios, significant differences in genetic variance were observed among scenarios (Additional File 11). Discrete genomic selection had significantly higher genetic variance than overlapping genomic selection, whereas discrete phenotypic selection led to significantly lower genetic variance than overlapping phenotypic selection (Additional File 12, 26).

### Selection accuracy

Significant differences in mean selection accuracy by scenario were observed in the RS-A cases (Additional File 11). Selection accuracy, as measured in the selected parents of the current generation per cycle, did not significantly differ between overlapping and discrete generations with replicated or unreplicated phenotypic selection (Additional File 12, 15—16). For selection on true genetic value, selection accuracy was by definition 1 for both discrete and overlapping generations. For genomic truncation selection, no differences in accuracy among overlapping and discrete generations were observed regardless of training set (Additional File 12, 18—19).

In genomic OCS, with the training set composed of all generations, selection accuracy was higher for overlapping generations in the short term if *N*_*e*_ = 100 and *h*^*2*^ = 0.5 (Additional File 12, 22). Overlapping generations also had higher accuracies in the medium term if *h*^*2*^ = 0.5 and *N*_*e*_ = (Additional File 12, 21). No significant differences were observed in the long term for OCS with training on all generations (Additional File 20—22). In genomic OCS with training on the previous five generations only, overlapping selection had higher selection accuracy in the short term only if *h*^*2*^ = 0.5 or 0.9 and *N*_*e*_ = 100 (Additional File 12, 25). In the medium term, overlapping selection had higher accuracies at all levels of *N*_*e*_ for *h*^*2*^ = 0.1, but only at *N*_*e*_ = 45 or 100 for *h*^*2*^ = 0.5 or 0.9 (Additional File 12, 23—25). In the long term, overlapping selection had higher accuracies at all levels of *h*^*2*^ and *N*_*e*_ observed with OCS and training on the previous five generations (Additional File 12, 23—25).

In the RS-AY cases, significant differences in mean selection accuracy were observed by scenario (Additional File 11). Discrete phenotypic selection produced higher selection accuracy than overlapping phenotypic selection, and discrete genomic selection produced higher selection accuracy than overlapping genomic selection (Additional File 12, 26).

### Mean parental age

By definition, the age of the selected parents under discrete generations was always one in the RS-A scenarios. Both thrice-replicated and unreplicated overlapping phenotypic truncation selection always resulted in mean parental age significantly greater than 1 for overlapping relative to discrete generations (Additional File 15—16, 28). Interestingly, selection on true genetic value always resulted in mean parental age significantly greater than 1 with overlapping generations in the long term, and in the medium term with *h*^*2*^ = 0.5 (Additional File 17, 28). With genomic truncation selection and training on all generations, mean parental age was always higher with overlapping generations (Additional File 18, 28). With truncation selection and training on the previous five generations, overlapping generations had significantly higher mean parental age except in the medium term at *h*^*2*^ = 0.1 (Additional File 19, 28). With genomic OCS and training on all generations, mean parental age in overlapping scenarios was not significantly different from discrete at *N*_*e*_ = 10 in the medium term only, but was significantly higher in the short and long terms (Additional File 20—22, 28). Mean parental age was always signficantly higher than discrete for *N*_*e*_ *=* 45 and 100 with genomic OCS and training on all generations (Additional File 21—22, 28). With genomic OCS and training on the previous five generations, mean parental age did not significantly differ between overlapping and discrete generations if *N*_*e*_ = 10 in the short term (Additional File 23, 28). However, at all other timepoints and levels of *N*_*e*_ overlapping selection led to significantly higher mean parental age than discrete (Additional File 23—25, 28).

In the RS-AY scenarios, mean parental age was 3.67 years under discrete selection. For the overlapping scenarios, mean parental age was significantly greater than 3.67 years with both phenotypic and genomic selection (Additional File 26—27).

## Discussion

The possibility of allowing generations to overlap in recurrent selection is not often considered. Although recycling a preferred parent across generations is common in applied breeding programs, nonpreferred individuals are generally discarded permanently. Here, the underlying theoretical basis for practicing discrete as opposed to overlapping recurrent phenotypic selection is demonstrated. Mean magnitude of error in selected individuals is larger than mean magnitude of error in the overall population, creating selection error bias. Over breeding cycles, selection error bias causes the magnitude of selection error to increase in phenotypically selected populations with overlapping generations. This propagation of selection error results in decreased genetic gain, whereas with discrete phenotypic selection the population recovers each cycle because the magnitude of the deviation of observed phenotypic value from true genetic value remains random in the selected individuals. Maintaining discrete generations in phenotypic selection prevents making the “same old mistakes” of selecting individuals erroneously believed to be exceptional repeatedly across cycles.

Notably, at higher heritabilities, the propagation of error takes more cycles to affect gain because the phenotypes of selected individuals deviate less from their true breeding value compared to at lower heritabilities. Discrete generations still outperformed overlapping generations if phenotypic observations were replicated three times, though the relative outperformance was slightly less than without replication as phenotypic value deviated less from true genetic value. However, with selection on true genetic value, no differences in mean genetic value were observed between discrete and overlapping generations, as is expected in absence of selection error.

The propagation of error under overlapping phenotypic selection can be thought of as failure to observe regression to a mean when individuals are not adequately evaluated; phenotypes at the tails of a distribution, far from the mean, are on average more likely to have larger magnitudes of error (Fig. 7). In breeding for population improvement, individuals in the upper tail of the phenotypic distribution—and outliers beyond the upper tail of the distribution— are inherently of interest. Many phenotypes are in the tails of the distribution due to error. In selection from discrete generations the total number of outliers is small, whereas in selection from overlapping generations the total number of outliers grows as breeding cycles are completed and total number of selection candidates grows. Thus, the number of highly erroneous phenotypes selected as parents is limited under discrete selection, and this restriction causes discrete phenotypic selection to outperform overlapping phenotypic selection. Though only three-fold replication of phenotypes is tested here, using additional replicates of phenotypic value should further restrict propagation of error in overlapping generations.

**Figure 7.**
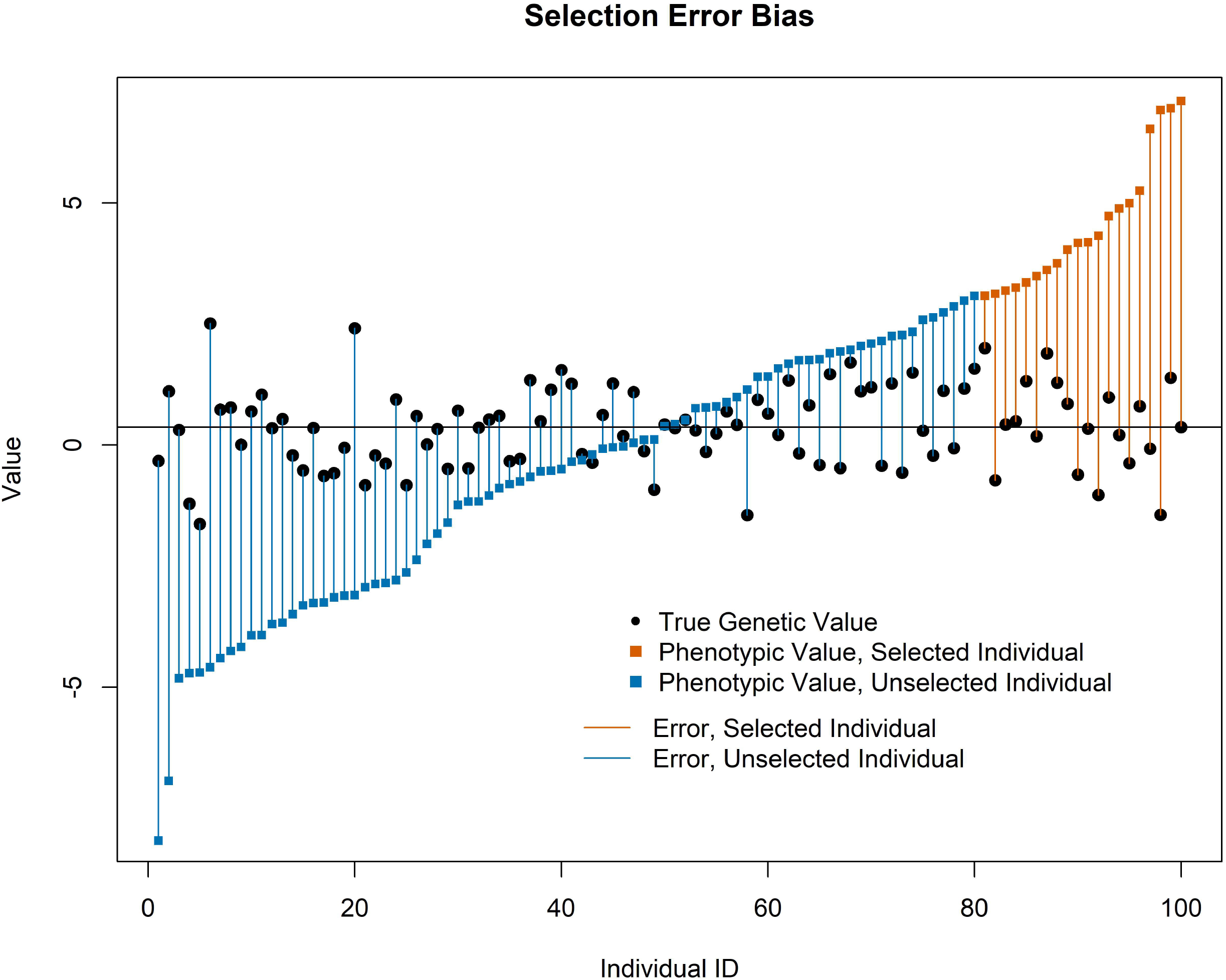
Selection error bias illustration. Phenotypic values, true genetic values, and errors of selected and unselected individual candidates at *h*^*2*^ = 0.1 in the first cycle of overlapping phenotypic selection for the RS-A pipeline. The magnitude of error is greater at the tails of the phenotypic values, including the upper tail from which individuals are selected.

The effect of overlapping vs. discrete generations in genomic truncation selection has not been previously evaluated to the authors’ knowledge. Mean genetic value does not significantly differ in discrete and overlapping genomic truncation selection, in contrast to phenotypic selection. Addition of new data to the model with each generation of genomic selection eliminates the problem of error propagation observed in phenotypic selection, as estimates of breeding value are improved by replicated observations of allele-phenotype combinations (which is synonymous with observations of more relatives). Though we hypothesized that overlapping generations might lead to more genetic gain than discrete as accuracy of GEBVs increased in older individuals with phenotyping of progeny, this was not the case due to the positive genetic trend from selection [18]. In other words, older individuals tended to have lower true genetic values than younger individuals in the presence of effective selection, so any increase in accuracy did not result in increased gain.Generally, the mean parental age did not substantially increase in overlapping genomic truncation selection compared to discrete (although the small increase observed was significant), indicating that parents with the best GEBVs were usually from the most recent generation or most recent past generations.

Because we observed in previous simulations that overlapping truncation selection underperformed discrete selection at high heritabilities in the long term due to inbreeding, we tested whether controlling genomic inbreeding by OCS led to greater mean genetic values in overlapping than discrete OCS scenarios. It is also well-established that genomic selection requires genomic control of inbreeding for maximal long-term gain, and at times genomic control of inbreeding can increase short-term gain relative to truncation selection [19-22]. However, we did not generally observe that overlapping selection outperformed discrete selection in OCS scenarios except at relatively high effective population size and high heritability. Interestingly, there is an explicit penalty to use of individuals from past generations in OCS due not to their genetic values but rather their addition to the rate of inbreeding [21]. If overlapping generations are allowed, control of inbreeding generally results from increasing the number of parents selected and not from increasing the generation interval in canonical OCS [18]. Thus, in contrast to genomic truncation selection, the relatively similar performance of overlapping and discrete OCS is likely due to the control of inbreeding as well as balance of gain per cycle and increased selection accuracy per cycle. With OCS at high *N*_*e*_ and *h*^*2*^ = 0.5 or 0.9, overlapping generations always had higher mean genetic values than discrete. This may indicate that overlapping generations allow more flexibility than discrete in balancing increases in inbreeding and genetic gain when inbreeding was more strictly constrained, as more individuals with more combinations of genetic value and relatedness were available to meet the constraints imposed. This is in agreement with the observation of Villanueva et al. (2000) that the optimal generation interval was higher with more stringent restrictions on inbreeding, as well as use of fewer parents [22].

As demonstrated in the RS-AY scenarios, error can propagate from any source with overlapping phenotypic selection— year error, genotype x year interaction error, or random plot error. Because we simulated greater plot error variance than year or genotype x year variance in stages from which parents were selected, we observed relatively more selection error bias due to plot error than other sources with overlapping phenotypic selection. Increasing the variance of the year or genotype x year values would likely increase their relative contributions to overall selection error; in applied breeding programs, the relative contribution of each source of error depends on the program. Additionally, we expect that selection error bias is not specific to plant breeding and can occur in other cyclical systems in which repeated selection occurs in the presence of random observational error.

The propagation of error was not restricted by movement of cohorts through advancement stages alone in the RS-AY scenario; restriction of propagation of error was accomplished by use of a statistical method to estimate breeding value. In the RS-AY scenarios, we only tested use of RR-BLUP to estimate breeding value. We expect that other estimation methods without a relationship matrix (e.g. best linear unbiased prediction) should restrict propagation of error if multi-year observations are available, as in the RS-AY scenario. However, if multiple observations are not available (as in the RS-A scenario), then estimation methods without relationship matrices would not restrict propagation of error.

To build on the conclusions of this study, it would be useful to test relative performance of overlapping and discrete generations under different genomic selection schemes, such as the modified reciprocal recurrent selection practiced in commercial hybrid breeding programs. Testing non-additive genetic architectures may also be relevant. Though speculative, it would also be interesting to test discrete and overlapping generations with multi-trait genomic selection. We hypothesize that in cases where multiple objectives are to be optimized (e.g. multiple phenotypic traits with different trait architectures), overlapping generations may provide more combinations of traits within genomic selection candidates and increase multi-trait gain.

## Conclusions

Based on the trends observed, generations should be kept discrete under recurrent mass phenotypic selection to avoid decreased genetic gain due to selection error bias. With genomic truncation selection, we observed no advantage to allowing overlapping generations under the assumptions used, though with genomic OCS it appeared the overlapping generations allowed more effective control of inbreeding than discrete generations at high effective population sizes with low targeted inbreeding rates.

## Declarations

### Ethics approval and consent to participate

Not applicable.

### Consent for publication

Not applicable.

### Availability of data and materials

All data generated or analysed during this study are included in this published article and its supplementary information files.

### Authors’ contributions

ML executed the study, wrote code for analysis, interpreted data, and drafted the manuscript. JR discovered selection error bias, wrote code for analysis, interpreted data, and edited the manuscript. Both authors read and approved the final manuscript.

## Acknowledgements

We thank Anthony J. Studer for advising and supporting ML throughout the course of this work. We thank Stephen P. Moose, Daniel Davidson, and David Slater for providing computational resources which enabled the study. We thank R. Chris Gaynor for developing AlphaSimR and providing code to model compound symmetry. We thank the anonymous reviewers of this work for their insights.

## Additional files

**Additional file 1**

Format: R programming language (.R)

Title: Script to generate base population

Description: R script used to generate the base population used in the study with the AlphaSimR package. Also contains custom optiSel functions used in the study.

**Additional file 2**

Format: R programming language (.R)

Title: Script to start RS-A simulations

Description: R script to initiate the RS-A simulations

**Additional file 3**

Format: R programming language (.R)

Title: Script to run RS-A simulations

Description: R script to run the RS-A simulations

**Additional file 4**

Format: R programming language (.R)

Title: Script to start RS-AY simulations

Description: R script to initiate the RS-AY simulations

**Additional file 5**

Format: R programming language (.R)

Title: Script to draw RS-AY year effects

Description: R script to save year effects for the RS-AY simulations

**Additional file 6**

Format: R programming language (.R)

Title: R script for RS-AY overlapping phenotypic selection scenario

Description: R script to run the simulation for the RS-AY phenotypic selection with overlapping generations scenario

**Additional file 7**

Format: R programming language (.R)

Title: R script for RS-AY discrete phenotypic selection scenario

Description: R script to run the simulation for the RS-AY phenotypic selection with discrete generations scenario

**Additional file 8**

Format: R programming language (.R)

Title: R script for RS-AY overlapping genomic selection scenario

Description: R script to run the simulation for the RS-AY genomic selection with overlapping generations scenario

**Additional file 9**

Format: R programming language (.R)

Title: R script for RS-AY discrete genomic selection scenario

Description: R script to run the simulation for the RS-AY genomic selection with discrete generations scenario

**Additional file 10**

Format: Microsoft Excel Workbook (.xlsx)

Title: Raw Simulation Results

Description: Excel file containing response values for all variable, cycles or years, and simulation replicates for the RS-A and RS-AY scenarios. See metadata tab for additional information.

**Additional file 11**

Format: Microsoft Word Document (.docx)

Title: Analyses of variance

Description: Results for all analyses of variance described in the study.

**Additional file 12**

Format: Microsoft Excel Workbook (.xlsx) Title: Contrasts

Description: Results for all contrasts described in the study.

**Additional file 13**

Format: Microsoft Word Document (.docx)

Title: RS-A Mean Genetic Values, Supplementary

Description: Plots of mean genetic value by cycle surrounded by 95% confidence intervals for the RS-A scenarios with genomic truncation selection and training on the previous five generations (fiveGen Trunc) as well as all RS-A OCS scenarios.

**Additional file 14**

Format: Microsoft Word Document (.docx)

Title: RS-A Selection Error Bias, Supplementary

Description: Plots of selection error bias by cycle surrounded by 95% confidence intervals for the RS-A scenarios with genomic truncation selection and training on the previous five generations (fiveGen Trunc) as well as all RS-A OCS scenarios.

**Additional file 15**

Format: Microsoft Word Document (.docx)

Title: RS-A Phenotypic Selection: All Responses

Description: Plots of all responses recorded for the RS-A phenotypic selection scenario.

**Additional file 16**

Format: Microsoft Word Document (.docx)

Title: RS-A Phenotypic 3rep Selection: All Responses

Description: Plots of all responses recorded for the RS-A phenotypic 3rep selection scenario.

**Additional file 17**

Format: Microsoft Word Document (.docx)

Title: RS-A True Genetic Value: All Responses

Description: Plots of all responses recorded for the RS-A true genetic value selection scenario.

**Additional file 18**

Format: Microsoft Word Document (.docx)

Title: RS-A allGen Trunc: All Responses

Description: Plots of all responses recorded for the RS-A genomic truncation selection with training on all previous generations scenario (allGen Trunc).

**Additional file 19**

Format: Microsoft Word Document (.docx)

Title: RS-A fiveGen Trunc: All Responses

Description: Plots of all responses recorded for the RS-A genomic truncation selection with training on the previous five generations scenario (fiveGen Trunc).

**Additional file 20**

Format: Microsoft Word Document (.docx)

Title: RS-A allGen OCS Ne = 10: All Responses

Description: Plots of all responses recorded for the RS-A genomic optimum contribution selection with training on all previous generations scenario at Ne = 10 (allGen OCS Ne = 10)

**Additional file 21**

Format: Microsoft Word Document (.docx)

Title: RS-A allGen OCS Ne = 45: All Responses

Description: Plots of all responses recorded for the RS-A genomic optimum contribution selection with training on all previous generations scenario at Ne = 45 (allGen OCS Ne = 45)

**Additional file 22**

Format: Microsoft Word Document (.docx)

Title: RS-A allGen OCS Ne = 100: All Responses

Description: Plots of all responses recorded for the RS-A genomic optimum contribution selection with training on all previous generations scenario at Ne = 100 (allGen OCS Ne = 100)

**Additional file 23**

Format: Microsoft Word Document (.docx)

Title: RS-A fiveGen OCS Ne = 10: All Responses

Description: Plots of all responses recorded for the RS-A genomic optimum contribution selection with training on the previous five generations scenario at Ne = 10 (fiveGen OCS Ne = 10)

**Additional file 24**

Format: Microsoft Word Document (.docx)

Title: RS-A fiveGen OCS Ne = 45: All Responses

Description: Plots of all responses recorded for the RS-A genomic optimum contribution selection with training on the previous five generations scenario at Ne = 45 (fiveGen OCS Ne = 45)

**Additional file 25**

Format: Microsoft Word Document (.docx)

Title: RS-A fiveGen OCS Ne = 100: All Responses

Description: Plots of all responses recorded for the RS-A genomic optimum contribution selection with training on the previous five generations scenario at Ne = 100 (fiveGen OCS Ne = 100)

**Additional file 26**

Format: Microsoft Word Document (.docx)

Title: RS-AY: All Responses

Description: Plots of all responses recorded for the RS-AY scenarios, including both phenotypic and genomic selection.

**Additional file 27**

Format: Microsoft Word Document (.docx)

Title: RS-AY Student’s *t*-tests

Description: Results of Student’s *t*-tests conducted for the RS-AY year error bias and mean parental age responses.

**Additional file 28**

Format: Microsoft Word Document (.docx)

Title: RS-A Student’s *t*-tests

Description: Results of Student’s *t*-tests conducted for the RS-A mean parental age responses.

## References

1. Duvick, D. N. (1996). Plant breeding, an evolutionary concept. Crop Science, 36(3), 539–548.

2. Harlan, J. R., De Wet, J. M. J., & Price, E. G. (1973). Comparative evolution of cereals. Evolution, 27(2), 311–325.

3. Hallauer, A. R., & Darrah, L. L. (1985). Compendium of recurrent selection methods and their application. Critical Reviews in Plant Sciences, 3(1), 1–33.

4. Eberhart, S. A. (1970). Factors effecting efficiencies of breeding methods. African soils, 15(1/3), 655–680.

5. Lee, E. A., & Tracy, W. F. (2009). Modern maize breeding. In Handbook of Maize (pp. 141–160). Springer, New York, NY.

6. Dudley, J. W. (2007). From means to QTL: The Illinois longLterm selection experiment as a case study in quantitative genetics. Crop Science, 47, S–20.

7. Lorenz, A. J., Chao, S., Asoro, F. G., Heffner, E. L., Hayashi, T., Iwata, H., … & Jannink, J. L. (2011). Genomic selection in plant breeding: knowledge and prospects. In Advances in agronomy (Vol. 110, pp. 77–123). Academic Press.

8. Goddard, M. E., & Hayes, B. J. (2007). Genomic selection. Journal of Animal breeding and Genetics, 124(6), 323–330.

9. Heffner, E. L., Sorrells, M. E., & Jannink, J. L. (2009). Genomic selection for crop improvement. Crop Science, 49(1), 1–12.

10. Jannink, J. L., Lorenz, A. J., & Iwata, H. (2010). Genomic selection in plant breeding: from theory to practice. Briefings in functional genomics, 9(2), 166–177.

11. Heslot, N., Jannink, J. L., & Sorrells, M. E. (2015). Perspectives for genomic selection applications and research in plants. Crop Science, 55(1), 1–12.

12. Faux, A. M., Gorjanc, G., Gaynor, R. C., Battagin, M., Edwards, S. M., Wilson, D. L., … & Hickey, J. M. (2016). AlphaSim: software for breeding program simulation. The plant genome, 9(3), 1–14.

13. Wellmann, R. (2019). Optimum contribution selection for animal breeding and conservation: the R package optiSel. BMC bioinformatics, 20(1), 1–13.

14. Gaynor, R. C., Gorjanc, G., Bentley, A. R., Ober, E. S., Howell, P., Jackson, R., … & Hickey, J. M. (2017). A twoLpart strategy for using genomic selection to develop inbred lines. Crop Science, 57(5), 2372–2386.

15. Pinheiro, J., Bates, D., DebRoy, S., & Sarkar, D. (2017). R Core Team (2017) nlme: linear and nonlinear mixed effects models. R package version 3. 1–131.

16. Lenth, R., Singmann, H., Love, J., Buerkner, P., & Herve, M. (2018). Emmeans: Estimated marginal means, aka least-squares means. R package version, 1(1), 3.

17. Hothorn, T., Bretz, F., Westfall, P., Heiberger, R. M., Schuetzenmeister, A., Scheibe, S., & Hothorn, M. T. (2016). Package ‘multcomp’. Simultaneous inference in general parametric models. Project for Statistical Computing, Vienna, Austria.

18. Villanueva, B., Bijma, P., & Woolliams, J. A. (2000). Optimal mass selection policies for schemes with overlapping generations and restricted inbreeding. Genetics Selection Evolution, 32(4), 1–17.

19. Meuwissen, T. H. E. (1997). Maximizing the response of selection with a predefined rate of inbreeding. Journal of animal science, 75(4), 934–940.

20. Jannink, J. L. (2010). Dynamics of long-term genomic selection. Genetics Selection Evolution, 42(1), 35.

21. Meuwissen, T. H. E., & Sonesson, A. K. (1998). Maximizing the response of selection with a predefined rate of inbreeding: overlapping generations. Journal of animal science, 76(10), 2575–2583.

22. Woolliams, J. A., Berg, P., Dagnachew, B. S., & Meuwissen, T. H. E. (2015). Genetic contributions and their optimization. Journal of Animal Breeding and Genetics, 132(2), 89–99.

