## Additional File 11 for "New cycle, same old mistakes? Overlapping vs. discrete generations in long-term recurrent selection"

**Table S2**

| **Predictor** | **Numerator *df*** | **Denominator *df*** | ***F*** | ***P*** |
| --- | --- | --- | --- | --- |
| *RS-A (Additive Trait)* | | | | |
| Mean genetic value | | | | |
| *Short term (cycle 5)* | | | | |
| Intercept | 1 | 588 | 137611.77 | < 0.0001 |
| Scenario | 65 | 588 | 169.49 | < 0.0001 |
| *Medium term (cycle 25)* | | | | |
| Intercept | 1 | 588 | 313395.2 | < 0.0001 |
| Scenario | 65 | 588 | 472.00 | < 0.0001 |
| *Long term (cycle 45)* | | | | |
| Intercept | 1 | 483 | 234555.82 | < 0.0001 |
| Scenario | 53 | 483 | 380.34 | < 0.0001 |
| Selection error bias | | | | |
| *Short term (cycle 5)* | | | | |
| Intercept | 1 | 588 | 28856.85 | < 0.0001 |
| Scenario | 65 | 588 | 365.32 | < 0.0001 |
| *Medium term (cycle 25)* | | | | |
| Intercept | 1 | 588 | 28934.82 | < 0.0001 |
| Scenario | 65 | 588 | 661.80 | < 0.0001 |
| *Long term (cycle 45)* | | | | |
| Intercept | 1 | 483 | 10686.08 | < 0.0001 |
| Scenario | 53 | 483 | 940.92 | < 0.0001 |
| Mean genomic inbreeding | | | | |
| *Short term (cycle 5)* |  |  |  |  |
| Intercept | 1 | 588 | 673887.40 | < 0.0001 |
| Scenario | 65 | 588 | 3076.90 | < 0.0001 |
| *Medium term (cycle 25)* |  |  |  |  |
| Intercept | 1 | 588 | 1203738.2 | < 0.0001 |
| Scenario | 65 | 588 | 11435.2 | < 0.0001 |
| *Long term (cycle 45)* |  |  |  |  |
| Intercept | 1 | 483 | 935270.7 | < 0.0001 |
| Scenario | 53 | 483 | 6337.1 | < 0.0001 |
| Genetic variance | | | | |
| *Short term (cycle 5)* | | | | |
| Intercept | 1 | 588 | 19654.15 | < 0.0001 |
| Scenario | 65 | 588 | 18.97 | < 0.0001 |
| *Medium term (cycle 25)* | | | | |
| Intercept | 1 | 588 | 6124.24 | < 0.0001 |
| Scenario | 65 | 588 | 70.24 | < 0.0001 |
| *Long term (cycle 45)* | | | | |
| Intercept | 1 | 483 | 449.26 | < 0.0001 |
| Scenario | 53 | 483 | 99.95 | < 0.0001 |
| Mean selection accuracy | | | | |
| *Short term (cycle 5)* | | | | |
| Intercept | 1 | 534 | 75745.08 | < 0.0001 |
| Scenario | 59 | 534 | 42.40 | < 0.0001 |
| *Medium term (cycle 25)* | | | | |
| Intercept | 1 | 534 | 141656.27 | < 0.0001 |
| Scenario | 59 | 534 | 71.24 | < 0.0001 |
| *Long term (cycle 45)* | | | | |
| Intercept | 1 | 362 | 101510.79 | < 0.0001 |
| Scenario | 47 | 362 | 108.61 | < 0.0001 |
| *RS-AY (Additive, Year, and Additive x Year Trait)* | | | | |
| Mean genetic value |  |  |  |  |
| Intercept | 1 | 36 | 19724 | < 0.0001 |
| Scenario | 3 | 36 | 14 | < 0.0001 |
| Selection error bias |  |  |  |  |
| Intercept | 1 | 36 | 4608 | < 0.0001 |
| Scenario | 3 | 36 | 204 | < 0.0001 |
| Plot error bias |  |  |  |  |
| Intercept | 1 | 36 | 2674 | < 0.0001 |
| Scenario | 3 | 36 | 109 | < 0.0001 |
| Additive x year error bias | | | | |
| Intercept | 1 | 36 | 1817 | < 0.0001 |
| Scenario | 3 | 36 | 3 | 0.0434 |
| Mean genomic inbreeding | | | | |
| Intercept | 1 | 36 | 1010 | < 0.0001 |
| Scenario | 3 | 36 | 37 | < 0.0001 |
| Genetic variance |  |  |  |  |
| Intercept | 1 | 36 | 420 | < 0.0001 |
| Scenario | 3 | 36 | 14 | < 0.0001 |
| Mean selection accuracy | | | | |
| Intercept | 1 | 36 | 86 | < 0.0001 |
| Scenario | 3 | 36 | 19 | < 0.0001 |

ANOVAs for the linear models used in the study.
