## Supplementary figures and images for "New cycle, same old mistakes? Overlapping vs. discrete generations in long-term recurrent selection"

### Additional File 13

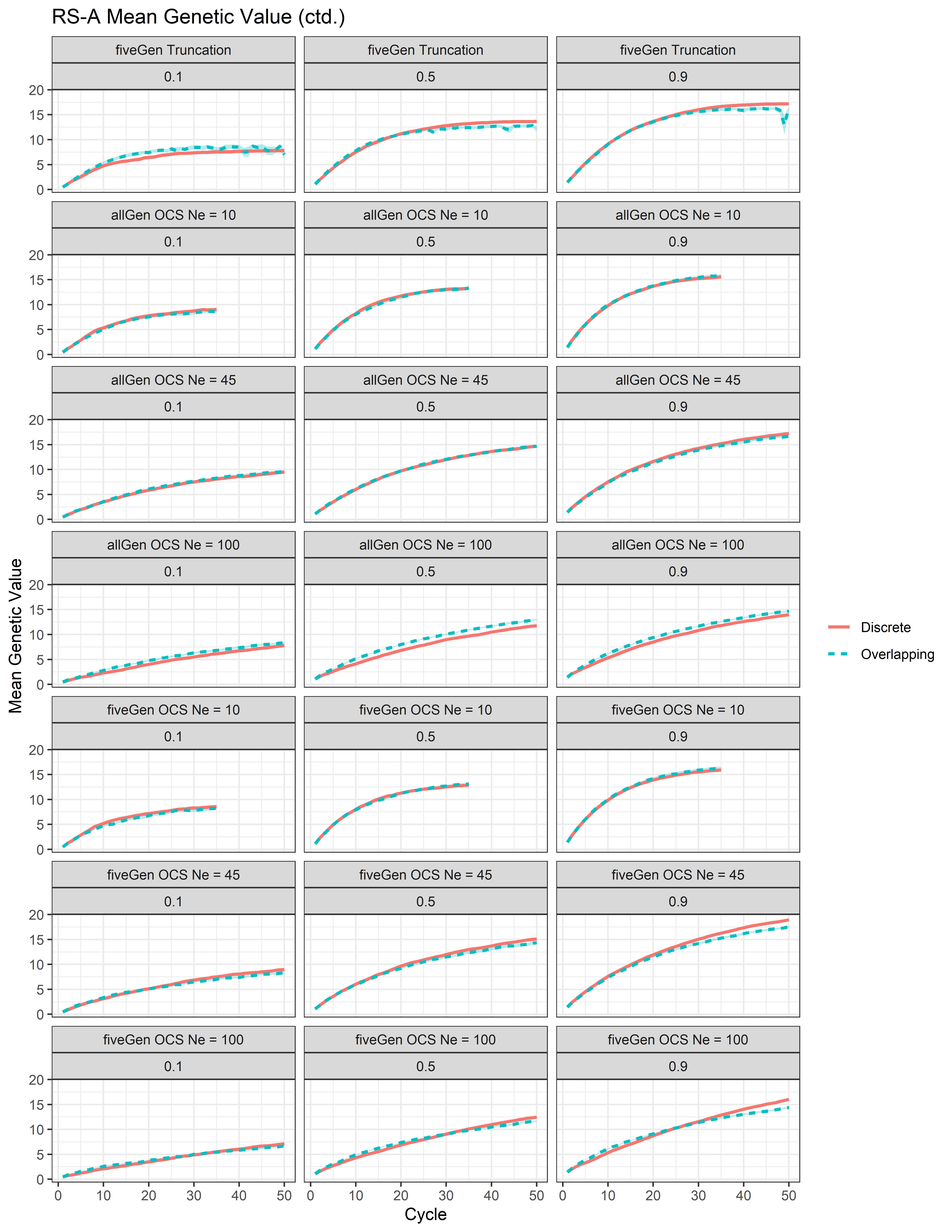

### Additional File 14

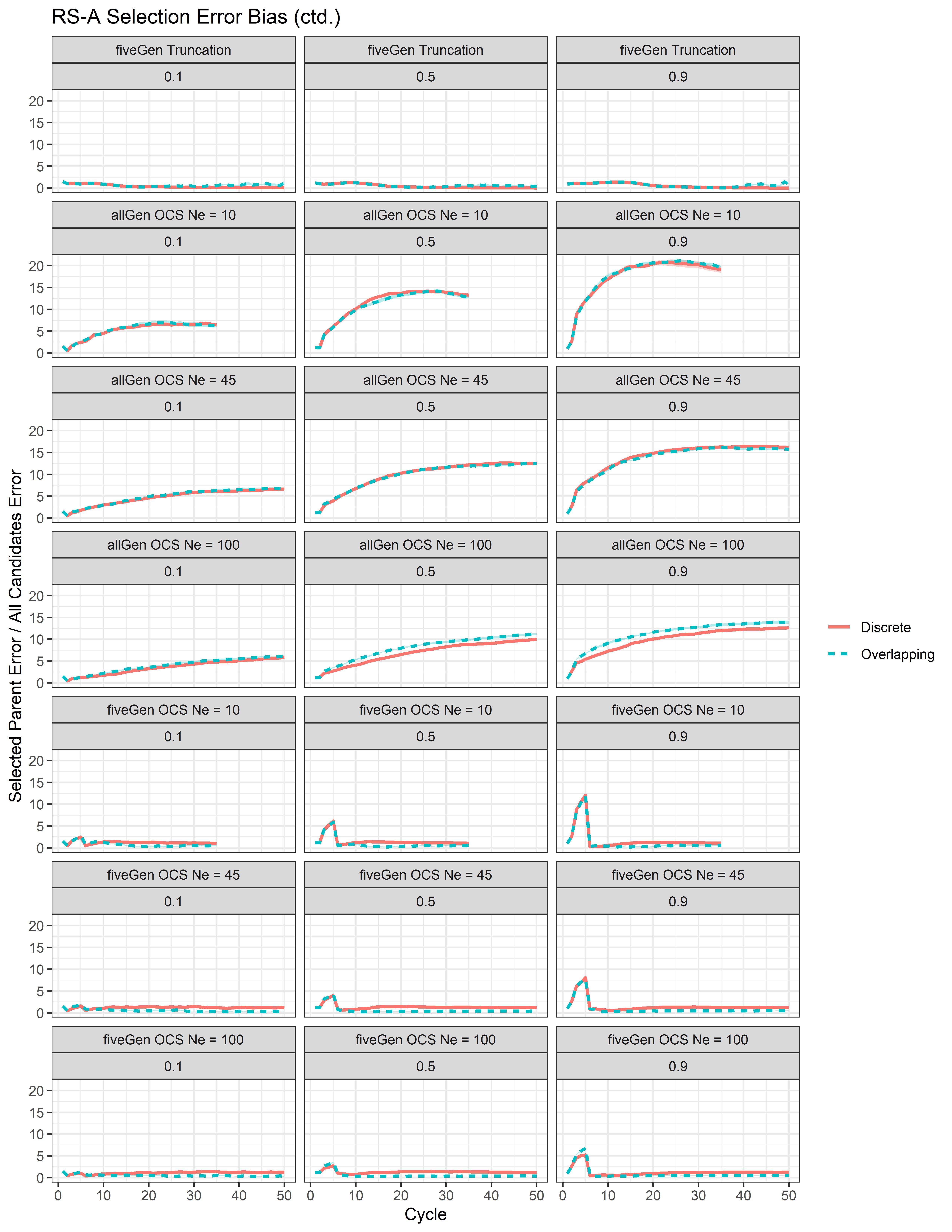

### Additional File 15

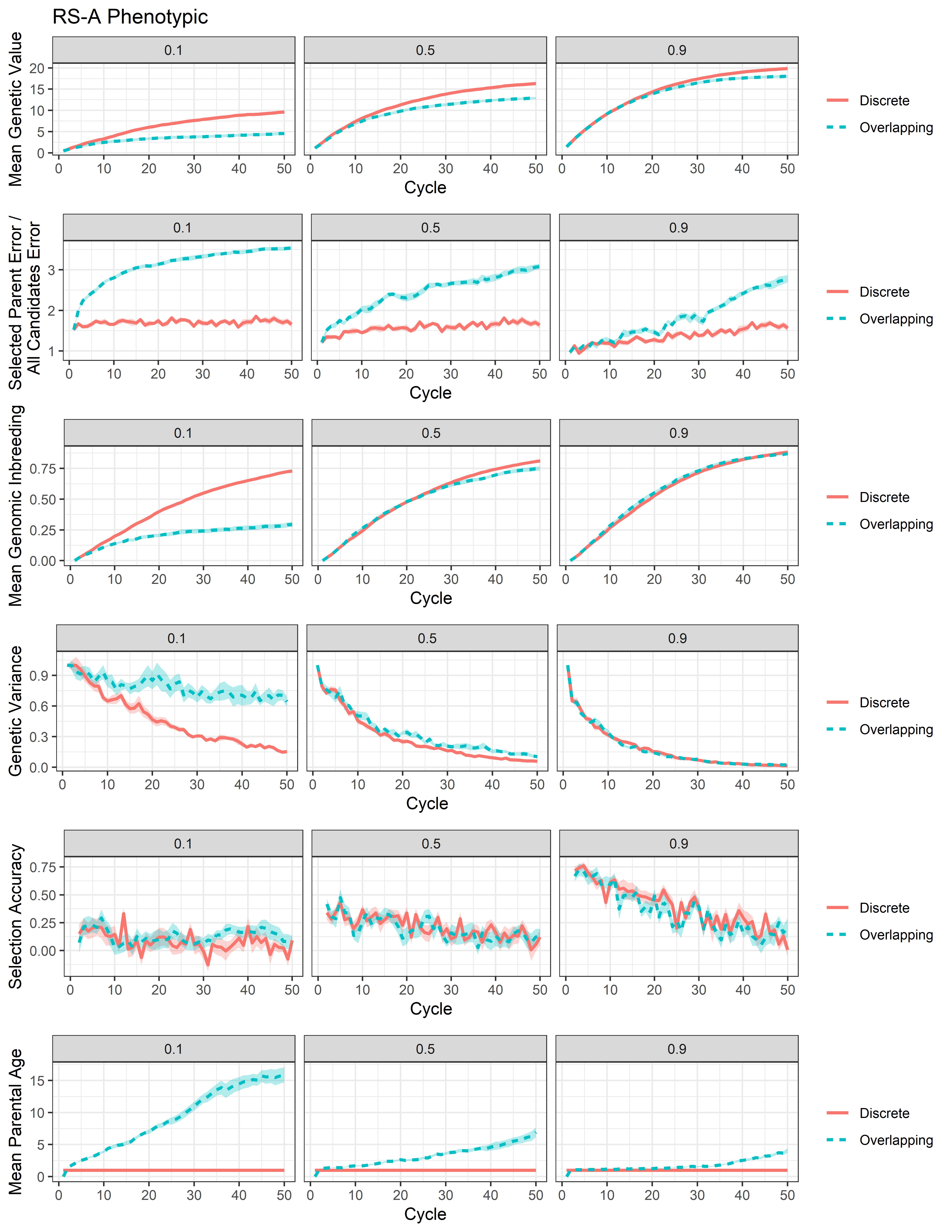

### Additional File 16

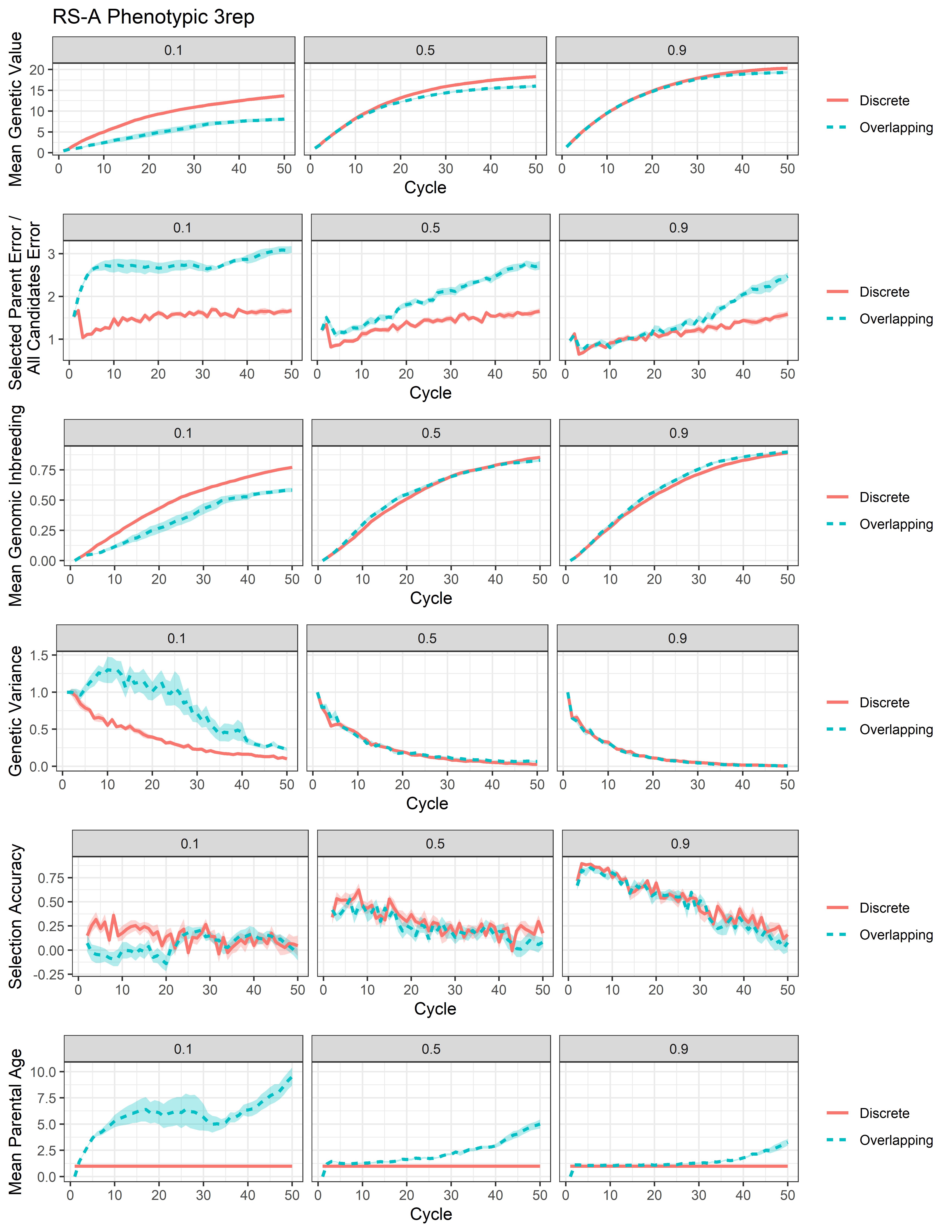

### Additional File 17

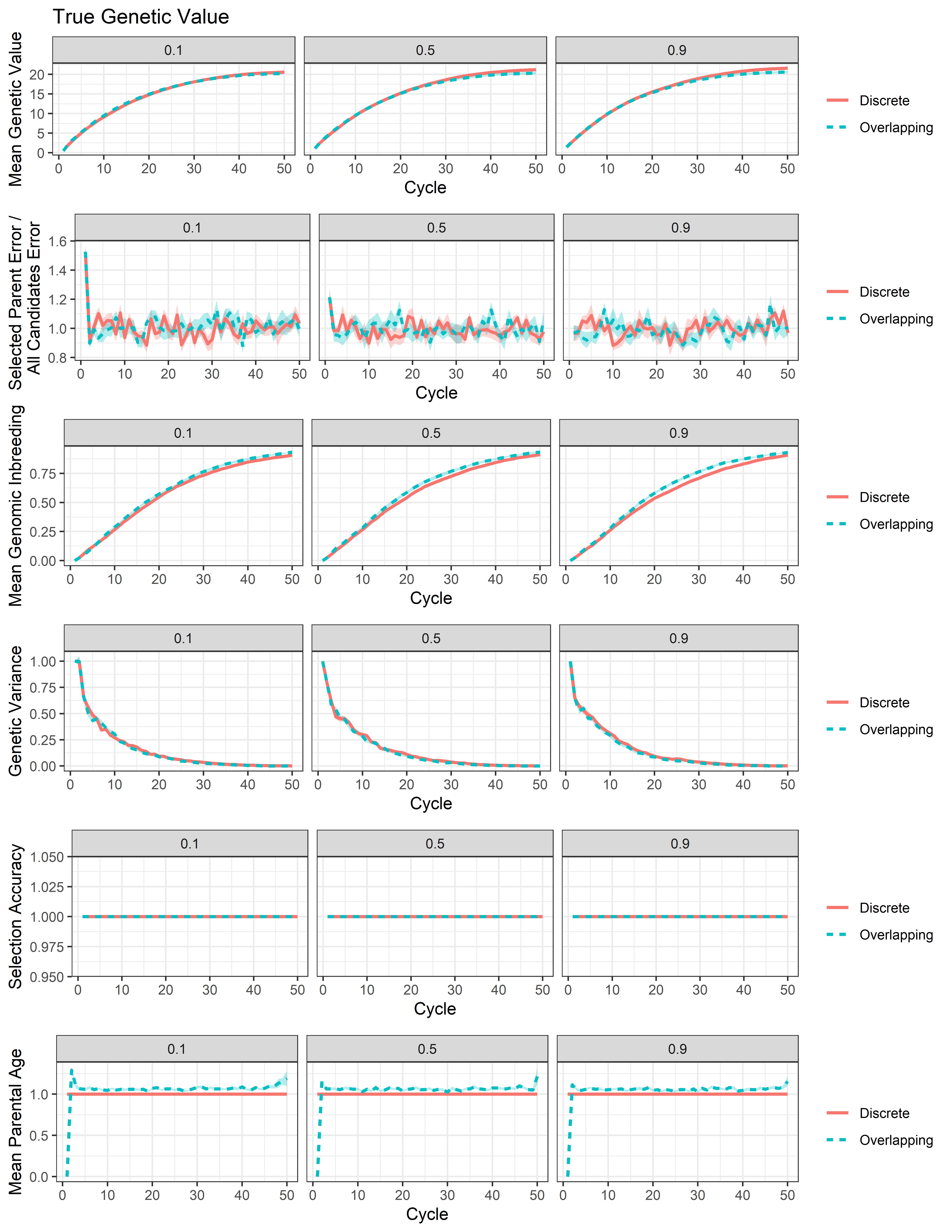

### Additional File 18

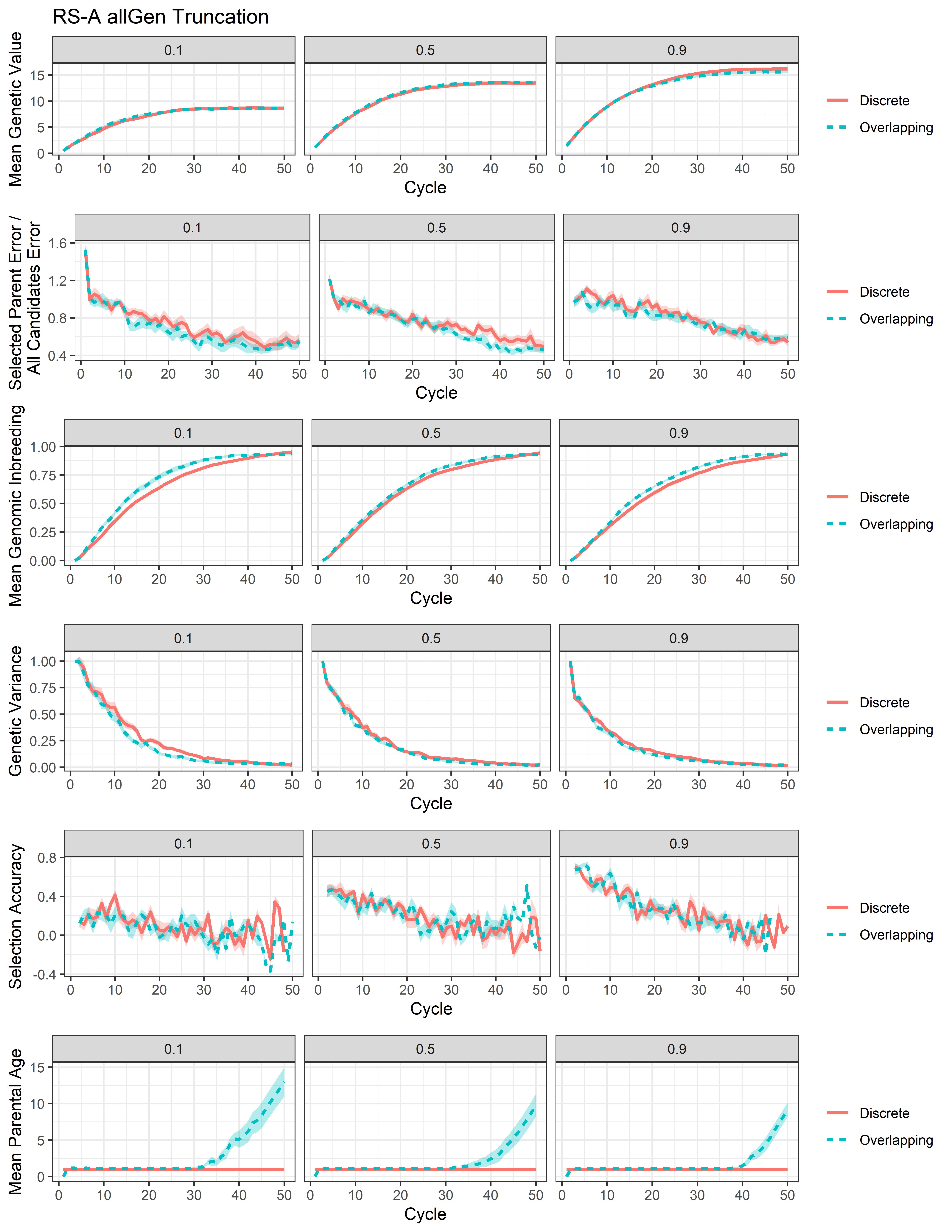

### Additional File 19

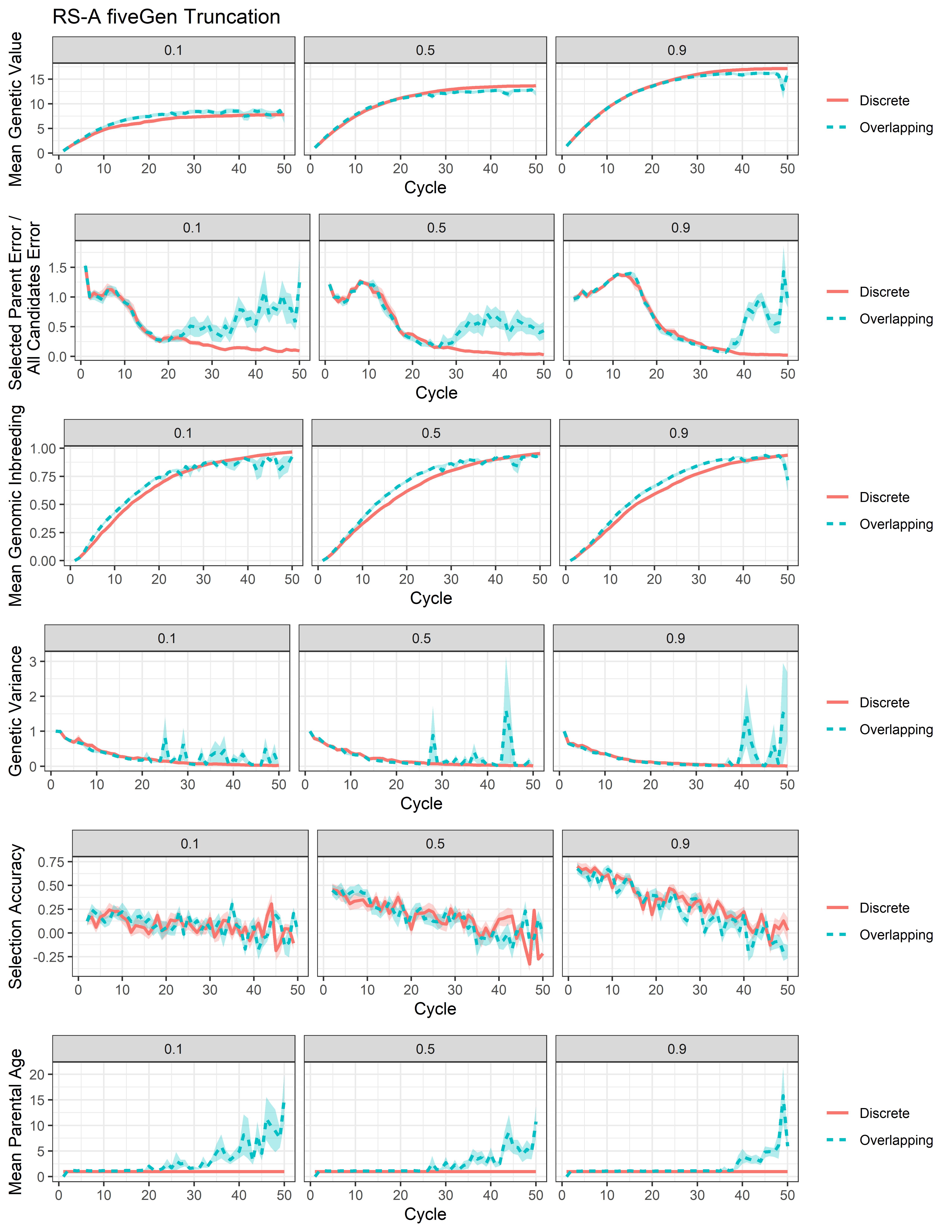

### Additional File 20

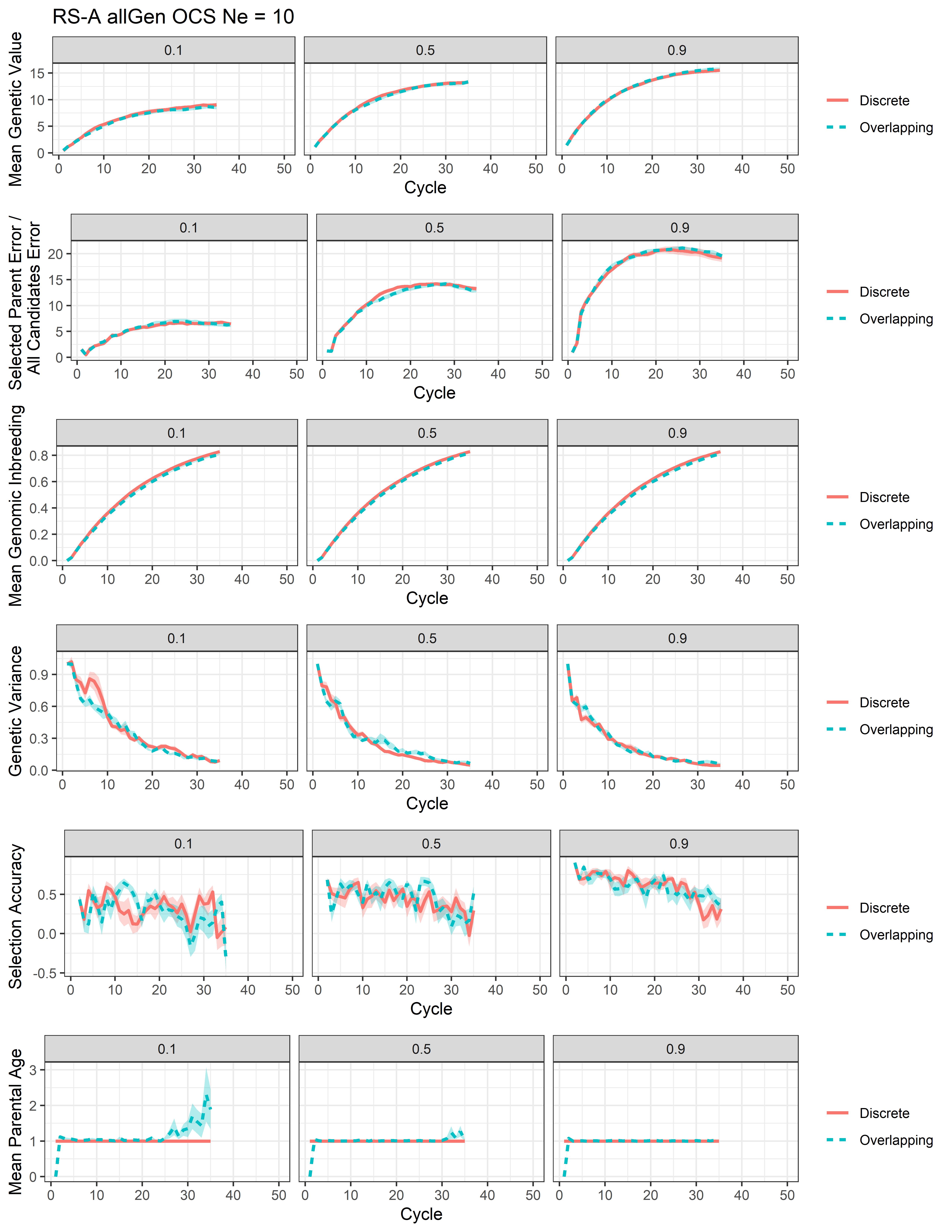

### Additional File 21

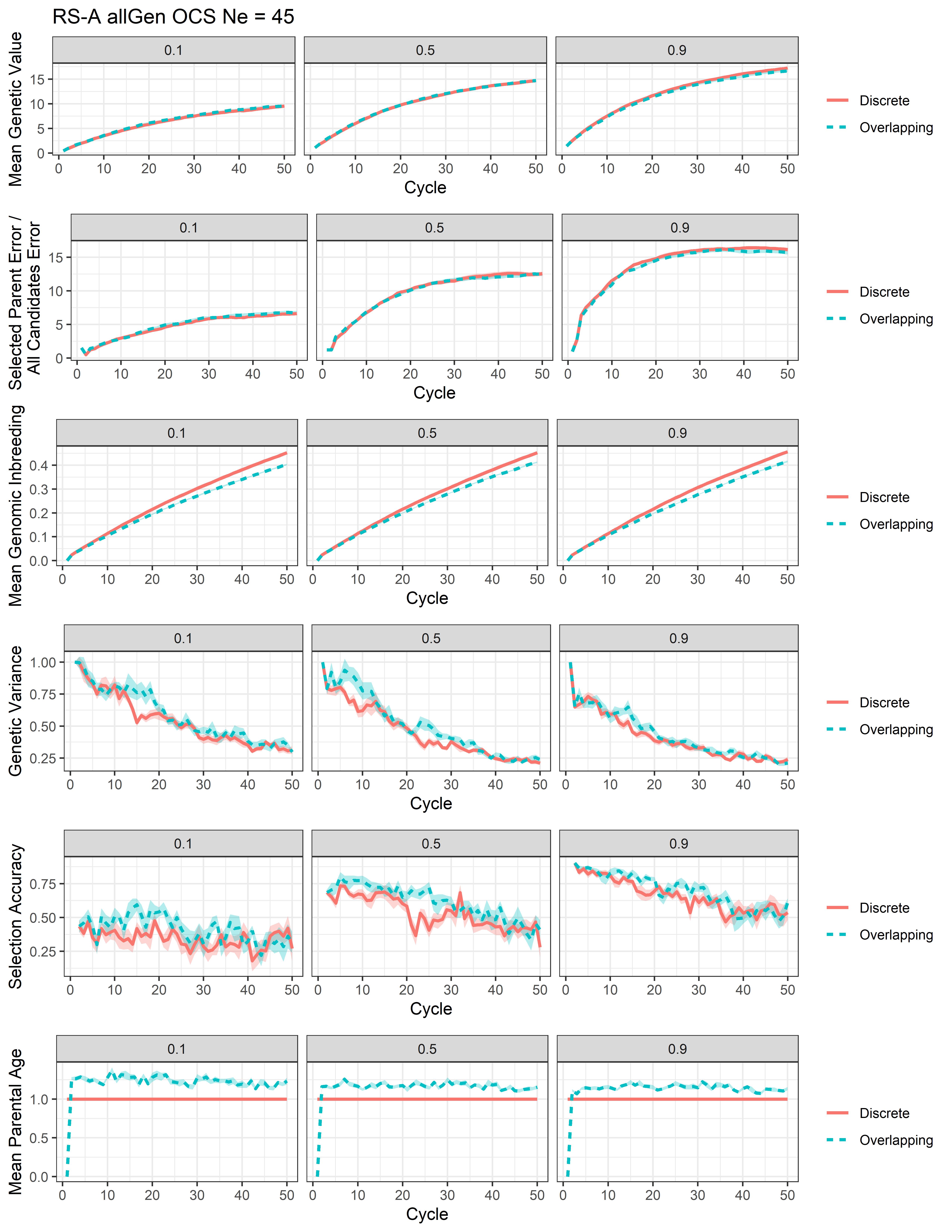

### Additional File 22

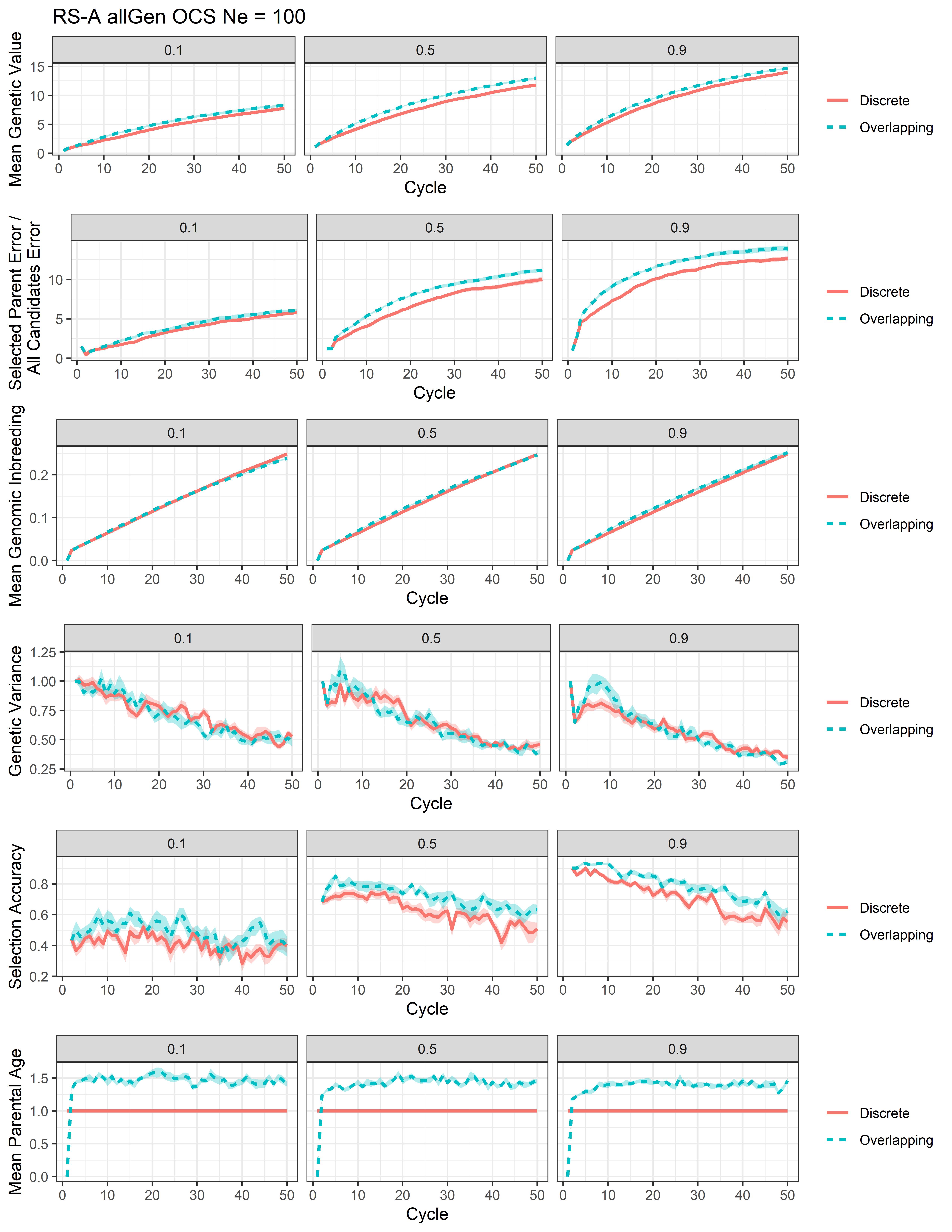

### Additional File 23

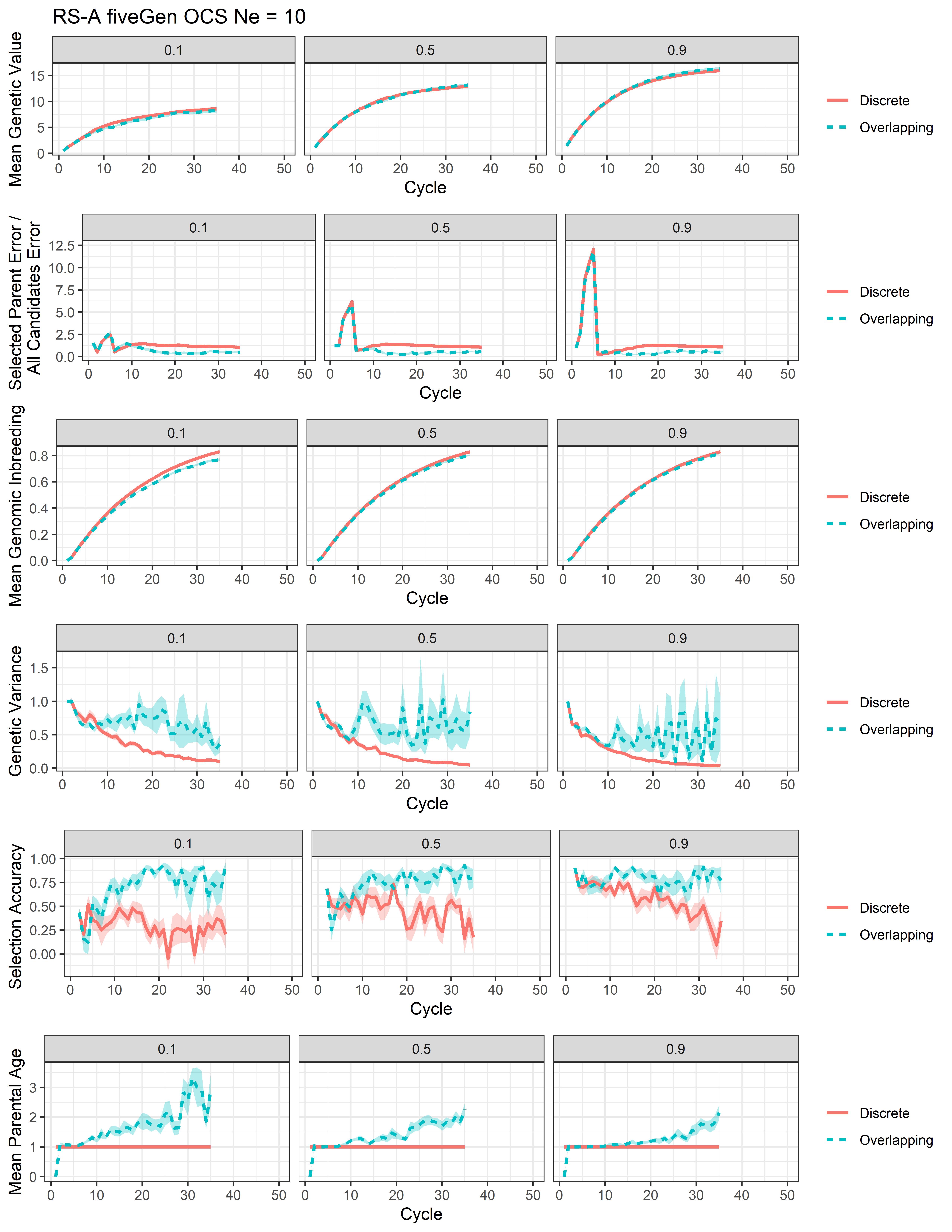

### Additional File 24

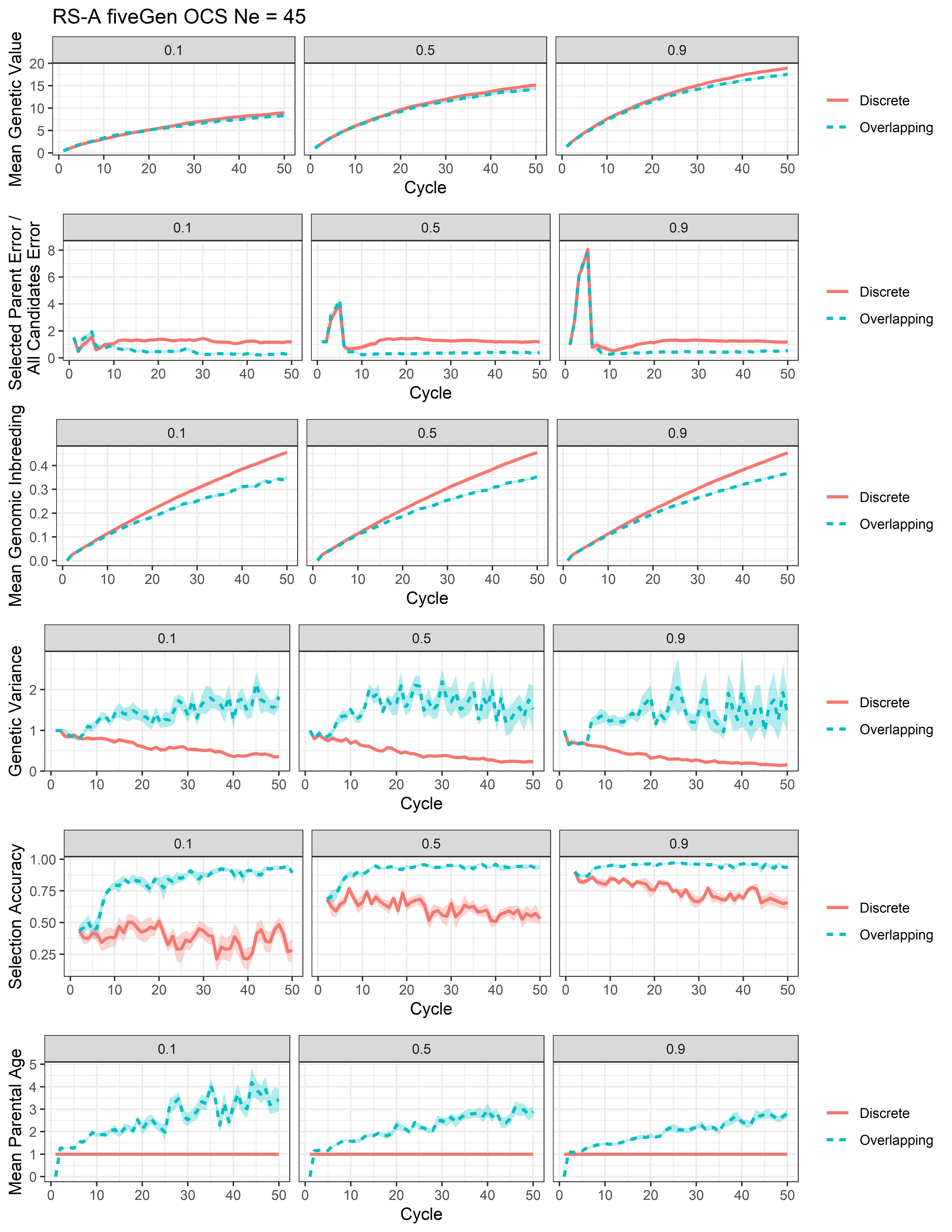

### Additional File 25

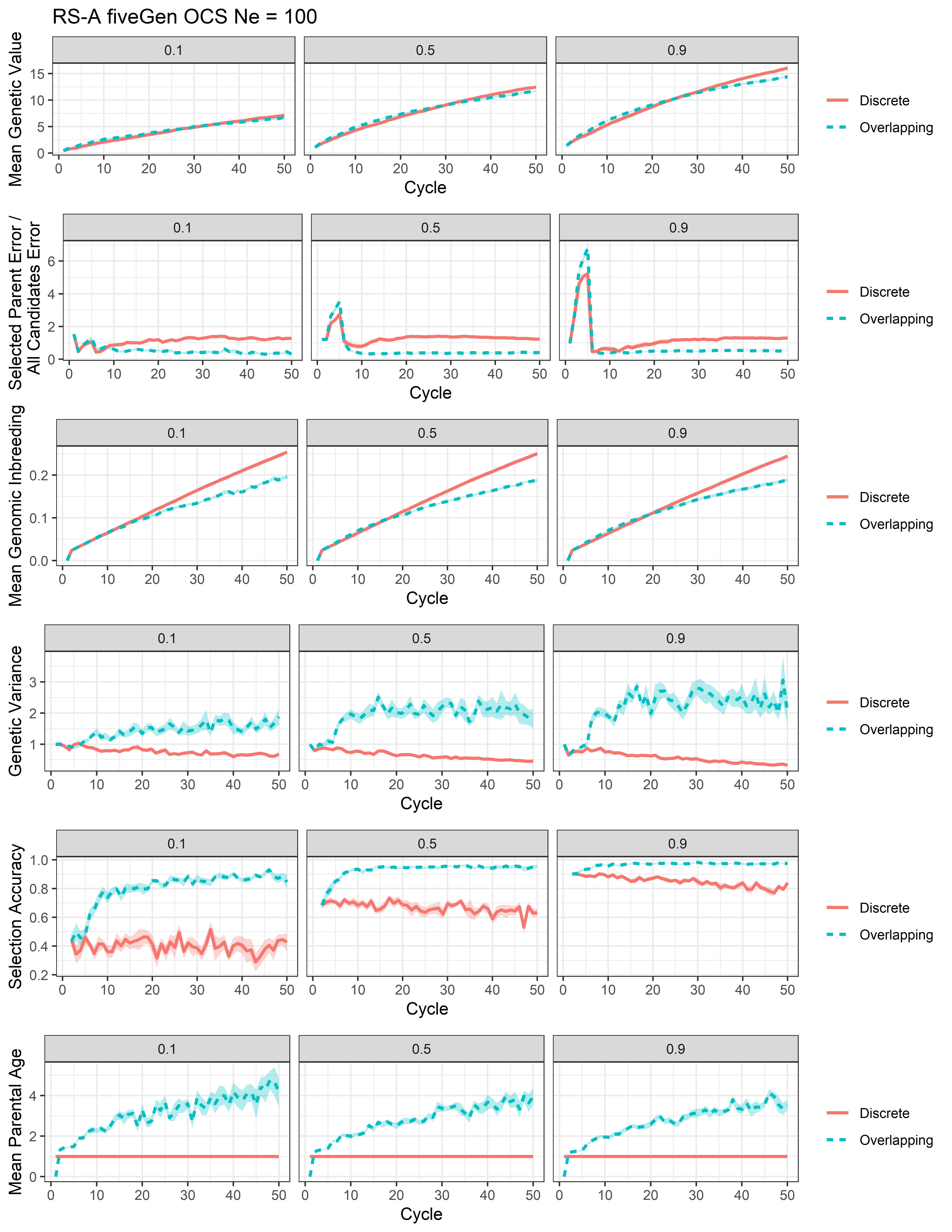

### Additional File 26

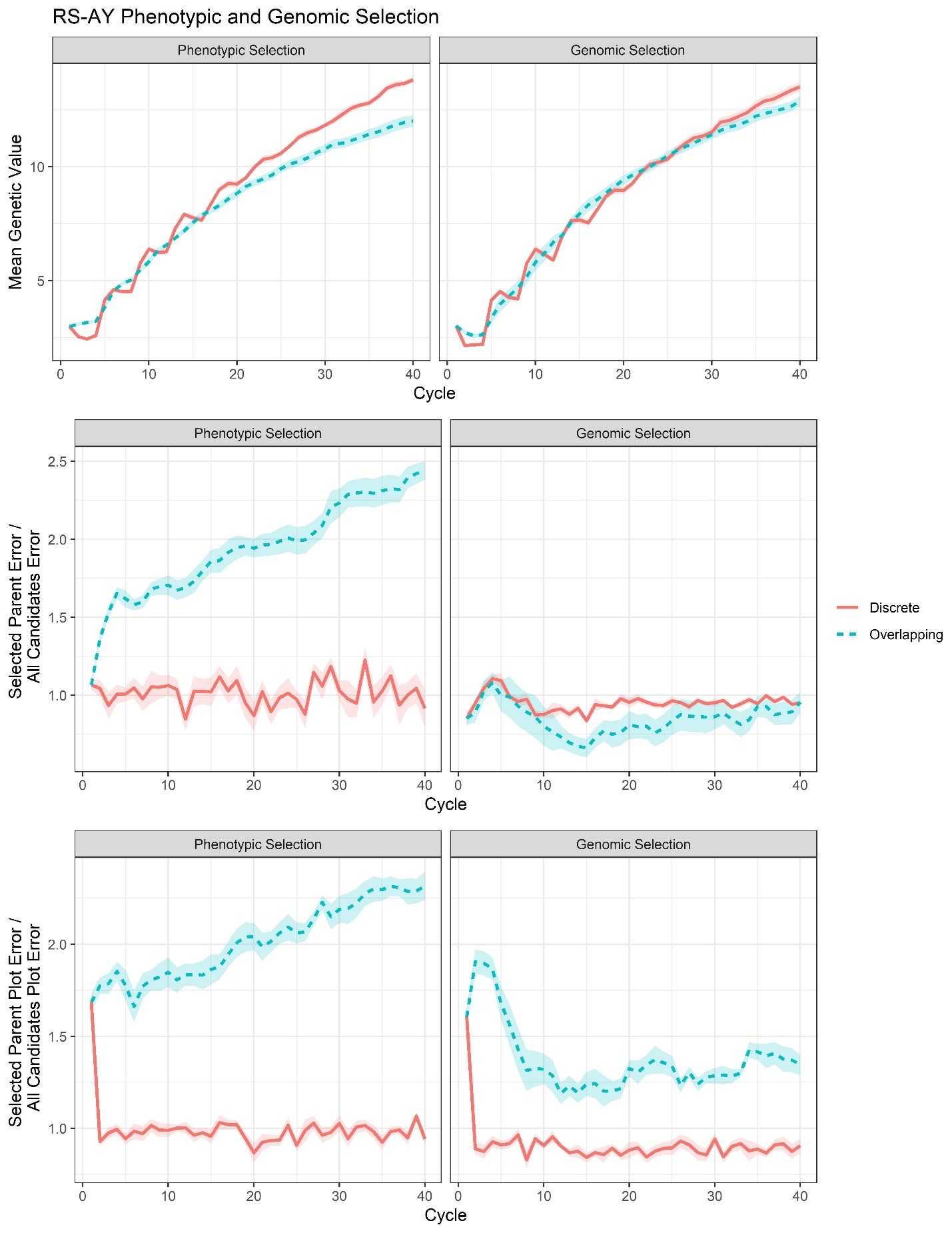

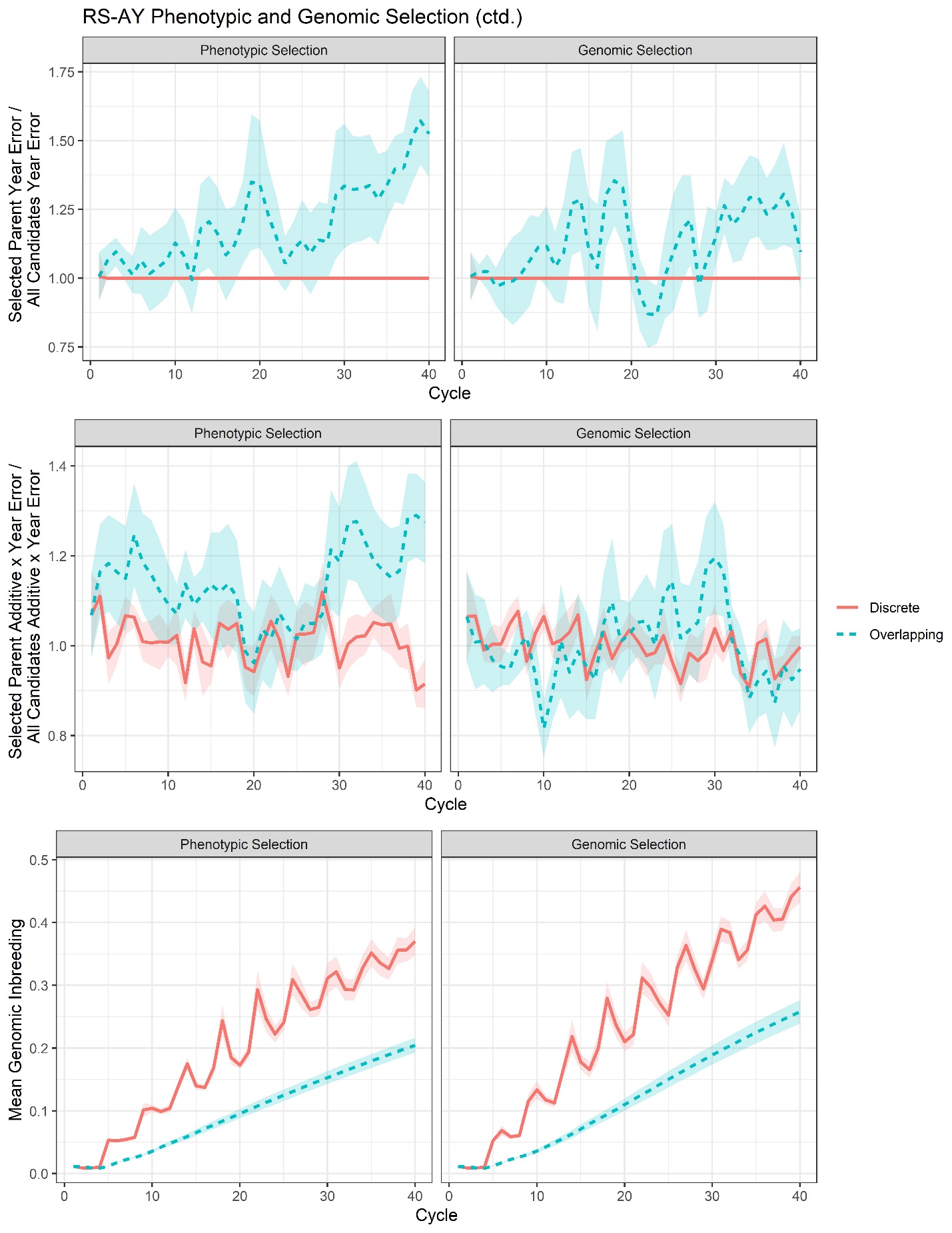

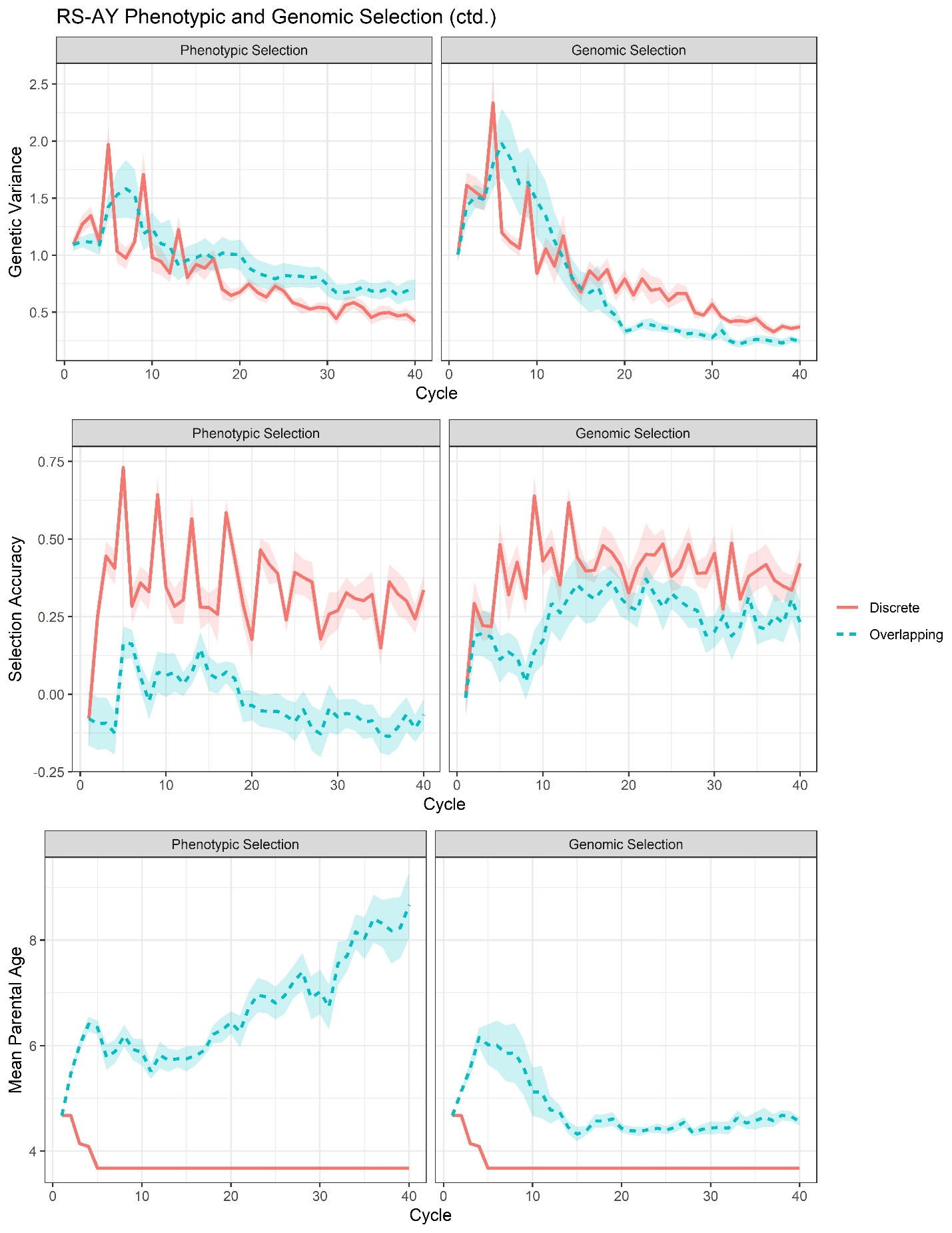
