## Additional File 27 for "New cycle, same old mistakes? Overlapping vs. discrete generations in long-term recurrent selection"

| **Scenario** | | ***t*** | | | **d.f.** | ***P*** |
| --- | --- | --- | --- | --- | --- | --- |
| Overlapping Phenotypic | | 8.071 | | | 9 | **< 0.0001** |
| Overlapping Genomic | | 12.451 | | | 9 | **< 0.0001** |

For the RS-AY scenarios, results of one-sample *t* tests of mean parental age at year 40 for each overlapped scenario compared to μ = 3.67. Means were compared to 3.67 because it was the average generation interval in years for the RS-AY scenario under discrete selection. The Bonferroni-corrected α value was 0.025.

| **Scenario** | ***t*** | **d.f.** | ***P*** |
| --- | --- | --- | --- |
| Overlapping Phenotypic | 3.589 | 9 | **0.006** |
| Overlapping Genomic | 0.919 | 9 | 0.382 |

For the RS-AY scenarios, results of one-sample *t* tests of mean year error bias at year 40 for each overlapped scenario compared to μ = 1. Means were compared to 1 because year error bias is by definition 1 under discrete selection. The Bonferroni-corrected α value was 0.025.
