## Additional File 28 for "New cycle, same old mistakes? Overlapping vs. discrete generations in long-term recurrent selection"

| **Scenario** | ***P*, Cycle 5** | ***P*, Cycle 25** | ***P*, Cycle 45** |
| --- | --- | --- | --- |
| allGen truncation, h^2^ = 0.1 | **< 0.0001** | **0.0002** | **0.0023** |
| fiveGen truncation, h^2^ = 0.1 | **0.0002** | 0.1026 | 0.0272 |
| allGen OCS Ne = 10, h^2^ = 0.1 | 0.2189 | 0.0482 |  |
| fiveGen OCS Ne = 10, h^2^ = 0.1 | 0.2189 | 0.0201 |  |
| allGen OCS Ne = 45, h^2^ = 0.1 | **< 0.0001** | **< 0.0001** | **0.0007** |
| fiveGen OCS Ne = 45, h^2^ = 0.1 | **< 0.0001** | 0.0229 | **< 0.0001** |
| allGen OCS Ne = 100, h^2^ = 0.1 | **< 0.0001** | **< 0.0001** | **0.0001** |
| fiveGen OCS Ne = 100, h^2^ = 0.1 | **< 0.0001** | **< 0.0001** | 0.0053 |
| Phenotypic, h^2^ = 0.1 | **< 0.0001** | **< 0.0001** | **< 0.0001** |
| Phenotypic, 3rep, h^2^ = 0.1 | **< 0.0001** | 0.0070 | **< 0.0001** |
| True Genetic Value, h^2^ = 0.1 | 0.0051 | 0.0037 | **0.0013** |
| allGen truncation, h^2^ = 0.5 | **< 0.0001** | **0.0003** | 0.0078 |
| fiveGen truncation, h^2^ = 0.5 | **0.0002** | **< 0.0001** | 0.0065 |
| allGen OCS Ne = 10, h^2^ = 0.5 | 0.1833 | 0.2744 |  |
| fiveGen OCS Ne = 10, h^2^ = 0.5 | 0.1833 | 0.0030 |  |
| allGen OCS Ne = 45, h^2^ = 0.5 | **0.0002** | **0.0003** | **0.0009** |
| fiveGen OCS Ne = 45, h^2^ = 0.5 | **0.0006** | **< 0.0001** | **0.0002** |
| allGen OCS Ne = 100, h^2^ = 0.5 | **< 0.0001** | **< 0.0001** | **< 0.0001** |
| fiveGen OCS Ne = 100, h^2^ = 0.5 | **< 0.0001** | **< 0.0001** | **< 0.0001** |
| Phenotypic, h^2^ = 0.5 | **0.0003** | **< 0.0001** | **< 0.0001** |
| Phenotypic, 3rep, h^2^ = 0.5 | **< 0.0001** | **< 0.0001** | **< 0.0001** |
| True Genetic Value, h^2^ = 0.5 | 0.0019 | **0.0011** | **0.0007** |
| allGen truncation, h^2^ = 0.9 | **0.0003** | 0.0046 | 0.0056 |
| fiveGen truncation, h^2^ = 0.9 | **< 0.0001** | 0.0078 | 0.0066 |
| allGen OCS Ne = 10, h^2^ = 0.9 | 0.1678 | 0.1696 |  |
| fiveGen OCS Ne = 10, h^2^ = 0.9 | 0.1678 | 0.0123 |  |
| allGen OCS Ne = 45, h^2^ = 0.9 | **0.0004** | **< 0.0001** | **< 0.0001** |
| fiveGen OCS Ne = 45, h^2^ = 0.9 | **0.0004** | **< 0.0001** | **< 0.0001** |
| allGen OCS Ne = 100, h^2^ = 0.9 | **< 0.0001** | **< 0.0001** | **0.0001** |
| fiveGen OCS Ne = 100, h^2^ = 0.9 | **< 0.0001** | **< 0.0001** | **< 0.0001** |
| Phenotypic, h^2^ = 0.9 | **0.0005** | **< 0.0001** | **< 0.0001** |
| Phenotypic, 3rep, h^2^ = 0.9 | 0.0119 | **0.0001** | **< 0.0001** |
| True Genetic Value, h^2^ = 0.9 | 0.0026 | 0.0095 | **< 0.0001** |

Results of one-sample *t* tests of mean parental age for each overlapped scenario compared to μ = 1 for discrete generations at cycles 5, 25, and 45 at Bonferroni-corrected α values of 0.0018, 0.0018, and 0.0024 respectively. Bolded values were statistically significant.
