## Additional File 1 (.R) for "New cycle, same old mistakes? Overlapping vs. discrete generations in long-term recurrent selection"

##################### load packages ############################

library(AlphaSimR)
library(abind)
library(pedigree)
library(optiSel)

###################### load internal or custom functions ############################

checkphen <- function (phen, columns, quiet, na.Sex = TRUE) 
{
  if (!is.data.frame(phen)) {
    stop("Argument phen must be a data frame.\n")
  }
  if ("data.table" %in% class(phen)) {
    phen <- as.data.frame(phen)
    setDF(phen)
  }
  missingColumns <- setdiff(columns, c(colnames(phen), "Breed", 
                                       "Sire", "Dam", "Sex"))
  if (length(missingColumns) > 0) {
    stop(paste0("The following columns are missing in phen: ", 
                paste(missingColumns, collapse = ", "), "\n"))
  }
  if ("Born" %in% columns) {
    if (!is.numeric(phen$Born)) {
      phen$Born <- as.numeric(as.character(phen$Born))
    }
    if (any(is.na(phen$Born)) && !quiet) {
      cat("There are NA-values in column 'Born'. These individuals can be \n selection candidates but are ignored for computing population means.\n")
    }
  }
  if ("Sex" %in% columns) {
    if (!("Sex" %in% colnames(phen))) {
      phen$Sex <- NA
    }
    if (!is.character(phen$Sex)) {
      phen$Sex <- as.character(phen$Sex)
    }
    if ((!na.Sex) && any(is.na(phen$Sex))) {
      stop("There are NA-values in column 'Sex'.\n")
    }
    if (!all(phen$Sex %in% c("male", "female", 
                             NA))) {
      stop("Some sexes are not coded as 'male' and 'female'.\n")
    }
    if (all(phen$Sex %in% c("male"))) {
      stop("Females are missing in data frame 'phen'.\n")
    }
    if (all(phen$Sex %in% c("female"))) {
      stop("Males are missing in data frame 'phen'.\n")
    }
  }
  if ("Indiv" %in% columns) {
    if (!is.character(phen$Indiv)) {
      phen$Indiv <- as.character(phen$Indiv)
    }
    if (any(is.na(phen$Indiv))) {
      stop("Some Individual IDs in column 'Indiv' are NA in argument phen.\n")
    }
    if (any(duplicated(phen$Indiv))) {
      stop("Some Individuals appear twice in column 'Indiv' of argument phen.\n")
    }
    if (!identical(rownames(phen), phen$Indiv)) {
      rownames(phen) <- phen$Indiv
    }
  }
  if ("Sire" %in% columns) {
    if (!("Sire" %in% colnames(phen))) {
      phen$Sire <- NA
      cat("Column 'Sire' was missing. It is set to NA.\n")
    }
    if (!is.character(phen$Sire)) {
      phen$Sire <- as.character(phen$Sire)
    }
  }
  if ("Dam" %in% columns) {
    if (!("Dam" %in% colnames(phen))) {
      phen$Dam <- NA
      cat("Column 'Dam' was missing. It is set to NA.\n")
    }
    if (!is.character(phen$Dam)) {
      phen$Dam <- as.character(phen$Dam)
    }
  }
  if ("Breed" %in% columns) {
    if (!("Breed" %in% colnames(phen))) {
      phen$Breed <- "missing"
    }
    if (!is.character(phen$Breed)) {
      phen$Breed <- as.character(phen$Breed)
    }
    if (any(is.na(phen$Breed))) {
      stop("Column 'Breed' of argument phen contains 'NA'.\n")
    }
    if ("unknown" %in% phen$Breed) {
      cat("Individuals with breed name 'unknown' are removed.\n")
      phen <- phen[phen$Breed != "unknown", ]
    }
  }
  phen
}


agecont.mod <- function(Pedig, use = rep(TRUE, times = nrow(Pedig)), maxAge = NA){
  Pedig <- checkphen(Pedig, columns = c("Indiv", "Born", "Sire", "Dam"), quiet = TRUE)
  Pedig$Born <- floor(Pedig$Born)
  Sire <- Pedig[use, "Sire"]
  Dam <- Pedig[use, "Dam"]
  ageM <- Pedig[use, "Born"] - Pedig[Sire, "Born"]
  ageF <- Pedig[use, "Born"] - Pedig[Dam, "Born"] 
  parents <- c(ageM, ageF) #parent ages when offspring was born
  parents <- parents[!is.na(parents) & parents >= 0]
  if(!is.na(maxAge)){
    parents <- parents[parents <= maxAge]
  }
  if(min(parents) == 0){
    cat("Individuals whose parents are born in the same year are removed.\n")
    parents <- parents[parents > 0]
  }
  q <- max(parents + 1, na.rm = TRUE)
  pOff <- data.frame(age = 1:q, parent = tabulate(parents + 1, nbins = q)/(length(parents)))
  L <- sum((pOff$age - 1)*pOff$parent)
  vp <- rev(cumsum(rev(pOff$parent[-1])))
  v <- data.frame(age = pOff$age, parent = c(vp, 0))
  data.frame(age = 1:q, male = 0.5*v$parent/L, female = 0.5*v$parent/L)
}

######################### make the starting environment ####################################

#create founder haplotypes
founderPop<-  runMacs(nInd=100, nChr=10, segSites=1000) #segsites is per chromosome!

#set simulation parameters
SP<- SimParam$new(founderPop)
SP$addTraitA(100) #add trait
SP$setSexes("no") 
SimParam$snpChips
SP$setTrackPed(FALSE)
SP$setTrackRec(TRUE)
SP$addSnpChip(50)

#set error variance (genetic variance= 1 by default)
#ve<- 4 #12-2019: replaced with variable heritabilities

save.image("SeedEnvironment.RData") #for each heritability, simrep, you should load the seed environment
