## Additional File 2 (.R) for "New cycle, same old mistakes? Overlapping vs. discrete generations in long-term recurrent selection"

##################### load packages ############################

library(AlphaSimR)
library(abind)
library(pedigree)
library(optiSel)

##################### load seed environment ############################
load("SeedEnvironment.RData")

##################### make the first year, common to all criteria, varying across simreps #######
##### to get the starting pop to vary across simreps, but keep the founderPop fixed, you have to create all seed environments in the same session #####
##### otherwise seed is set and they will be identical ####

SP$setVarE(h2 = 0.1, varE = 4)
#SP$setVarE(h2 = 0.5, varE = 4) #use this line for h2 = 0.5
#SP$setVarE(h2 = 0.9, varE = 4) #use this line for h2 = 0.9

#results arrays that will be populated
#was: row is year, column is criteria, depth is simrep; was c(year, criteria, heritability, simulation)
#is: year, criteria (sim and heritability set externally)
RsltMat<- array(dim = c(50, 42)) #genetic gains
SdRsltMat <- array(dim = c(50, 42)) #sd on genetic gains
AgeMat<- array(dim = c(50, 42)) #ages
RsMat<- array(dim = c(50, 42)) #residuals
RsMatTrue<- array(dim = c(50, 42)) #true residuals
ErMatP<- array(dim = c(50, 42)) #error, parents
ErMatAl<- array(dim = c(50, 42)) #error, all
AccMat <- array(dim = c(50, 42)) #accuracies, all
AccPMat <- array(dim = c(50, 42)) #parent accuracies
AccTMat <- array(dim = c(50, 42)) #training set accuracies
AccCMat <- array(dim = c(50, 42)) #breeding candidate accuracies
#MeanInbrMatP <- array(dim = c(50, 42)) #parent inbreeding (pedigree)
#MeanGInbrMatP <- array(dim = c(50, 42)) #parent inbreeding (genomic)
#SdInbrMatP <- array(dim = c(50, 42)) #parent inbreeding sd (pedigree)
#SdGInbrMatP <- array(dim = c(50, 42)) #parent inbreeding sd (genomic)
#MeanInbrMatAl <- array(dim = c(50, 42)) #current generation inbreeding (pedigree)
MeanGInbrMatAl <- array(dim = c(50, 42)) #current generation inbreeding (genomic)
#SdInbrMatAl <- array(dim = c(50, 42)) #current generation inbreeding sd (pedigree)
SdGInbrMatAl <- array(dim = c(50, 42)) #current generation inbreeding sd (genomic)
#Fispx0Mat <- array(dim = c(50,10,20)) #Fis parent against pop0
#Fiscx0Mat <- array(dim = c(50,10,20)) #Fis curgen against pop0
#Fispx1Mat <- array(dim = c(50,10,20)) #Fis parent against p1
#Fiscx1Mat <- array(dim = c(50,10,20)) #Fis curgen against p1
MatAll_VarG <- array(dim = c(50, 42)) #genetic variance all
MatPar_VarG <- array(dim = c(50, 42)) #genetic variance parents
MatCur_VarG <- array(dim = c(50, 42)) #genetic variance curgen
MatAll_VarP <- array(dim = c(50, 42)) #phenotypic variance all
MatPar_VarP <- array(dim = c(50, 42)) #phenotypic variance parents
MatCur_VarP <- array(dim = c(50, 42)) #phenotypic variance current gen
#SolveMat <- array(dim = c(50, 3, 42))


############## start the simulation #####################
#save results for each year
mns <- array(dim = c(50)) #holds mean genetic value
sdmns <- array(dim = c(50))
psel <- c() #holds the selected genotypes
errsP <- array(dim = c(50))
errsAll <- array(dim = c(50))
Rs <- array(dim = c(50))
avgAge <- array(dim = c(50))
RsTrue <- array(dim = c(50))
acc <- array(dim = c(50))
accpar <- array(dim = c(50))
acctrain <- array(dim = c(50))
acccur <- array(dim = c(50))
#minbr_par <- array(dim = c(50))
#minbr_cur <- array(dim = c(50))
#mginbr_par <- array(dim = c(50,18))
mginbr_cur <- array(dim = c(50))
#sinbr_par <- array(dim = c(50))
#sinbr_cur <- array(dim = c(50))
#sginbr_par <- array(dim = c(50,18))
sginbr_cur <- array(dim = c(50))
#Fis_px0 <- matrix(nrow = 50, ncol = 18)
#Fis_cx0 <- matrix(nrow = 50, ncol = 18)
#Fis_px1 <- matrix(nrow = 50, ncol = 18)
#Fis_cx1 <- matrix(nrow = 50, ncol = 18)
varg_allgen <- array(dim = c(50))
varg_pargen <- array(dim = c(50))
varg_curgen <- array(dim = c(50))
varp_allgen <- array(dim = c(50))
varp_pargen <- array(dim = c(50))
varp_curgen <- array(dim = c(50))
#solvecheck <- array(dim = c(50, 3))
#colnames(solvecheck) <- c("valid", "solver", "status")

#make the starting pop, which is always the same (seed set)
pop0 <- newPop(founderPop) 
pop0 <- setPheno(pop=pop0) #the phenotypes change every time this is run

idsel <- pop0@id[order(pop0@pheno, decreasing=T)][1:20] #take the 20 best phenotypes as parents

p1 <- randCross(pop0, nCrosses=100, parents=match(idsel, pop0@id)) #new pop is born from the 20 best chosen (current gen)

allPops <- pop0 #merge pop0 with allPops table
#end founder pop creation
#save the first year info
varg_curgen[1] <- varG(pop0)[1,]
varp_curgen[1] <- varP(pop0)[1,]
varg_pargen[1] <- varG(pop0[idsel])[1,]
varp_pargen[1] <- varP(pop0[idsel])[1,]
errsP[1] <- mean(c(abs(pop0@pheno[match(idsel, pop0@id)]-pop0@gv[match(idsel, pop0@id)]))) #errors for the selected inds
errsAll[1] <- mean(c(abs(pop0@pheno-pop0@gv))) #mean error for all
mginbr_cur[1] <- 0
sginbr_cur[1] <- 0
mns[1] <- meanG(p1) #track the mean genetic values of the current gen
sdmns[1] <- 0
avgAge[1] <- 0 #we know the average age is 0
Rs[1] <- 0 #no ebvs; we assume the mean gv is 0
RsTrue[1] <- mean(pop0@gv[match(idsel, pop0@id)])-mean(pop0@gv) #gain in selected inds vs mean
varg_allgen[1] <- varG(allPops)[1,]
varp_allgen[1] <- varP(allPops)[1,] #phenotype lags genotype by 1 cycle

#for the subsequent 50 or so years, make up the phenotypes and run the regime
#when we start here:
#pop0 is the starting population, made from base
#p1 is the result of crossing phenotypically selected parents from pop0 (year 1)
#p is the current generation
#allPops is founder + current, but no phenos on current
#critPops is founder + current in the loop

p1 <- setPheno(pop=p1) #set phenotypes for the result of the randCross


##save.image("h1SeedEnvironment1.RData")
##save.image("h1SeedEnvironment2.RData")
##save.image("h1SeedEnvironment3.RData")
##save.image("h1SeedEnvironment4.RData")
##save.image("h1SeedEnvironment5.RData")
##save.image("h1SeedEnvironment6.RData")
##save.image("h1SeedEnvironment7.RData")
##save.image("h1SeedEnvironment8.RData")
##save.image("h1SeedEnvironment9.RData")
##save.image("h1SeedEnvironment10.RData")
