## Additional File 4 (.R) for "New cycle, same old mistakes? Overlapping vs. discrete generations in long-term recurrent selection"

library(AlphaSimR)
library(sommer)
'%notin%' <- Negate('%in%')


##### Function to make the compound symmetry phenotypes #####
## Creates a phenotype for 1 year and 1 replicate
# Adjust VarPlot to get right heritability
csphen <- function(pop, yearEff, VarPlot, VarGxY, VarGen){
  genEff = gv(pop)
  plotEff = rnorm(nInd(pop), sd=sqrt(VarPlot))
  
  # Create GxY effects
  Q = pullQtlGeno(pop) - 1 # QTL genotypes coded as -1,0,1
  
  addEffVar = popVar(matrix(SP$traits[[1]]@addEff))[1]
  gxyMarkerEffVar = addEffVar*VarGxY/VarGen # Variance for GxY marker effects
  gxyMarkerEff = rnorm(ncol(Q), sd=sqrt(gxyMarkerEffVar)) # Sample GxY marker effects
  
  gxyEff = Q%*%gxyMarkerEff
  
  popVar(gxyEff) # approximately VarGxY
  
  pheno = cbind(genEff + rep(yearEff, times = nInd(pop)) + gxyEff + plotEff,
                genEff, rep(yearEff, times = nInd(pop)), gxyEff, plotEff)
  
  return(pheno)
}
  


##### SIMULATION PARAMETERS ----
VarGen = 1
VarYear = 0.2
VarGxY = 0.2


#create founder haplotypes
founderPop <- runMacs(nInd=100, nChr=10, segSites=1000) #segsites is per chromosome!

#set simulation parameters
SP <- SimParam$new(founderPop)
SP$addTraitA(nQtlPerChr = 100, var = VarGen)
SP$setSexes("no")
SP$setTrackPed(FALSE)
SP$setTrackRec(TRUE)
SP$addSnpChip(50)

#reload for each rep to avoid setTrackRec getting too big across reps
#save.image(file = "RealisticStart.RData")
load("RealisticStart.RData")

for(REP in 10){ #change this number to make each of 10 reps
################################################################################  
# Fill the pipeline
################################################################################
  
  # Pull the initial parents
  Parents <- newPop(founderPop)
  ParentsPhen <- csphen(Parents, rnorm(1, sd=sqrt(VarYear)), 3.6, VarGxY, VarGen)
  Parents@pheno[,1] <- ParentsPhen[,1]
  Parents@fixEff <- rep(as.integer(0), times = nInd(Parents))
  
  # Presample year p-value for genotype-by-year interactions
  # These values are only used for filling the pipeline
  #P = runif(7)
  
  # start saving ages
  bY <- data.frame(ind = NA, year = NA)
  bY <- rbind(cbind(Parents@id, rep(0, times = length(Parents@id))))
  
  for(cycle in 1:7){
    print(paste("cycle", cycle))
    yearEff <- rnorm(1, sd=sqrt(VarYear)) #sets the year effect
    
    #year 1: make 100 random biparental crosses
    # Crossing is performed to produce 100 biparental populations. Parental combinations are chosen from all possible combinations
    # for the 50 parental lines in the crossing block (1225 possible combinations) using random sampling without replacement
    F1 <- randCross(Parents, nCrosses = 100, nProgeny = 97)
    bY <- rbind(bY, cbind(F1@id, rep(cycle, times = length(F1@id)))) #save birth years

    
    # year 2: make 97 DH lines per cross (not 100- controlled cost)
    # 97 doubled haploid lines are produced from each biparental family.
    if(cycle < 7){
    DH <- makeDH(F1, nDH = 1, simParam = SP)
    }
    bY <- rbind(bY, cbind(DH@id, rep(cycle, times = length(DH@id))))
    
    #year 3: make headrows from the DH lines (100 reps per DH) and select best 500 lines
    # The newly developed doubled haploids are planted in headrows to increase seed and perform visual selection. Visual selection
    # in the headrows is modeled as selection on a yield phenotype with heritability of 0.1 to represent the breeder selecting on
    # correlated traits. The breeder advances 500 lines.
    if(cycle < 6){
      Headrow <- DH
      HeadrowPhen <- csphen(Headrow, yearEff, 7.6, VarGxY, VarGen)
      Headrow@pheno[,1] <- HeadrowPhen[,1]
      Headrow@fixEff <- rep(as.integer(cycle), times = nInd(Headrow))
      Headrow <- selectInd(Headrow, nInd = 500)
    }
    
    
    #year 4: run a preliminary yield trial for the selected lines, then select 50 for advance, 20 to recycle to year 1
    # The 500 lines are evaluated in the PYT. The PYT represents evaluation in an unreplicated trial that is mechanically
    # harvested to measure yield. Selection in the PYT is modeled as selection on a yield phenotype with heritability of 0.2.
    # The best performing 50 lines are advanced to the next trial. The best performing 20 lines are also advanced to the next
    # year's crossing block, thereby completing a crossing cycle.
    if(cycle < 5){
      PYT <- Headrow
      PYTPhen <- csphen(Headrow, yearEff, 3.6, VarGxY, VarGen)
      PYT@pheno[,1] <- PYTPhen[,1]
      PYT@fixEff <- rep(as.integer(cycle), times = nInd(PYT))
      PYT <- selectInd(PYT, nInd = 50)
    }
    
    #year 5: run a small MET for the selected lines. Select 10 for advance. Same 10 go to crossing block.
    # The 50 lines advanced from the PYT are evaluated in an advanced yield trial (AYT). 
    # The AYT represents evaluation in a small, multilocation replicated yield trial. 
    # Selection in the AYT is modeled as selection on a yield phenotype with heritability of
    # 0.5. This value of heritability was based on the assumption that an AYT represents four effective
    # replications of the PYT. The best performing 10 lines in the AYT are advanced to the next
    # trial. These 10 lines are also considered as candidates for next year's crossing block. 
    # The next year's crossing block of 50 lines is composed of the 20 best PYT lines, the 10 best AYT lines,
    # and the best 20 lines selected from the current crossing block's 30 non-PYT lines.
    if(cycle < 4){
      AYT <- PYT
      AYTPhen <- csphen(AYT, yearEff, 0.6, VarGxY, VarGen)
      AYT@pheno[,1] <- AYTPhen[,1]
      AYT@fixEff <- rep(as.integer(cycle), times = nInd(AYT))
      AYT <- selectInd(AYT, nInd = 10)
    }
    
    #year 6: conduct elite yield trial for selected lines, advance all; update crossing block phenos
    # The 10 advanced lines are evaluated in an elite yield trial (EYT).
    # The EYT represents evaluation in a large, multilocation replicated yield trial. 
    # Selection in the EYT is modeled as selection on a yield phenotype with heritability of 0.67. This value of
    # heritability was based on the assumption that an EYT represents eight effective replications of the PYT. 
    # All 10 lines are kept in the EYT to be reevaluated in the following year. Any of
    # the 10 lines used in the current year's crossing block have their
    # phenotypes updated to reflect their performance in this year's
    # EYT before lines are chosen for next year's crossing block.
    if(cycle < 3){
      EYT <- AYT
      EYTPhen <- csphen(EYT, yearEff, 0.1, VarGxY, VarGen)
      EYT@pheno[,1] <- EYTPhen[,1]
      EYT@fixEff <- rep(as.integer(cycle), times = nInd(EYT))
    }
    
    #year 7: reevaluate eyt lines, update crossing block phenos
    # The 10 lines from the previous year's EYT are reevaluated. Any
    # of those still in the current crossing block have their phenotypes
    # updated to reflect their performance in both years of evaluation
    # in the EYT before lines are chosen for next year's crossing block.
    if(cycle < 2){
      EYT2 <- EYT
      EYT2Phen <- csphen(EYT2, yearEff, 0.1, VarGxY, VarGen)
      EYT2@pheno[,1] <- EYT2Phen[,1]
      EYT2@fixEff <- rep(as.integer(cycle), times = nInd(EYT2))
    }
    
    #year 8: variety
    # The line with the best average performance over the previous 2
    # yr of EYT evaluation is released as a variety.
    VT <- EYT2
    VT@pheno <- (EYT@pheno + EYT2@pheno) / 2
    VT <- selectInd(VT, nInd = 1)
  }
  save.image(file = paste("STARTENV", REP, ".RData", sep = ""))
}
  
  
###############################################################################
# End filling pipeline. Begin breeding program.
##############################################################################
